# Closed-loop robotic interactions reveal dynamic social filtering in schooling fish

**DOI:** 10.64898/2026.08.03.742553

**Authors:** Vaios Papaspyros, Yacine Barhoumi, Ramón Escobedo, Francesco Mondada, Clément Sire, Guy Theraulaz

## Abstract

Collective motion emerges from local interactions among individuals, yet whether interaction rules inferred from trajectory data correspond to the mechanisms actually used by animals remains unresolved. Here, we address this question using an autonomous closed-loop robotic fish implementing a data-driven model of social interactions reconstructed from the schooling fish *Hemigrammus rhodostomus*. The robot continuously updated its behavior from real-time tracking of freely swimming fish while reproducing spontaneous locomotion, wall avoidance, and anisotropic attraction and alignment. We compared entirely biological groups, biohybrid groups containing one robotic fish, and numerical simulations using identical behavioral descriptors across isolated individuals, pairs, and groups of five fish. The robotic fish successfully integrated into natural schools and reproduced the principal signatures of collective coordination, providing the first direct causal validation of interaction rules reconstructed from behavioral trajectories. Biohybrid experiments showed that a robot responding only to its single most influential neighbor was sufficient to sustain natural collective coordination. By contrast, numerical simulations reproduced the behavior of biological groups most accurately when each fish interacted with its two most influential neighbors. This discrepancy identifies the contribution of hydrodynamic interactions, which remain available to living fish but are absent from the robotic controller, demonstrating that physical and behavioral interactions jointly shape collective organization. These findings establish closed-loop biohybrid robotics as a powerful framework for experimentally testing the mechanisms underlying collective animal behavior.

**Significance:** Inferring the behavioral mechanisms underlying collective animal behavior from trajectory data alone cannot establish causality. We combined a data-driven model of fish social interactions with an autonomous closed-loop robotic fish that continuously interacted with freely swimming conspecifics. This biohybrid approach provides the first direct causal validation of interaction rules reconstructed from behavioral trajectories. Comparing biological groups, biohybrid groups, and numerical simulations further reveals that hydrodynamic interactions complement social interactions in shaping collective organization. While living fish require information from their two most influential neighbors to reproduce natural schools, a robotic fish lacking hydrodynamic feedback achieves comparable coordination by responding to only its single most influential neighbor, demonstrating the power of closed-loop biohybrid robotics for testing mechanisms of collective behavior.

## 1 Introduction

Coordinated collective behavior represents one of the most striking examples of distributed biological intelligence. Fish schools, bird flocks, insect swarms and many other animal societies achieve remarkable levels of coordination without centralized control, relying instead on the continuous exchange of locally acquired social information among interacting individuals Vicsek and Zafeiris [2012]. Through this decentralized organization, groups collectively detect predators Treherne and Foster [1981], Ward et al. [2011], navigate through complex environments Grünbaum [1998], locate resources and maintain cohesion while remaining highly responsive to rapidly changing conditions Sumpter [2010], Xue et al. [2023], Lin et al. [2025]. Understanding how these collective capabilities emerge from local interactions has consequently become a central question connecting behavioral ecology, neuroscience, statistical physics, robotics and artificial intelligence Giardina [2008], Lopez et al. [2012], Herbert-Read [2016], Ouellette [2022].

During the past four decades, research on collective behavior has progressively shifted from describing emergent spatial patterns toward understanding the informational processes that generate them. Early theoretical studies demonstrated that simple attraction, repulsion and alignment rules were sufficient to reproduce many characteristics of schools and flocks Aoki [1982], Okubo [1986], Huth and Wissel [1992, 1994], Niwa [1994], Couzin et al. [2002]. These models established self-organization as a fundamental principle of collective behavior, but they did not determine whether living animals actually implement the proposed interaction rules. As behavioral tracking became increasingly quantitative, interaction functions began to be reconstructed directly from experimental trajectories Katz et al. [2011], Herbert-Read et al. [2011], Gautrais et al. [2012], Calovi et al. [2018]. This transition transformed attraction and alignment from theoretical assumptions into experimentally measurable behavioral functions.

Recent work suggests, however, that these interaction functions represent only the observable outcome of a deeper computational process. Rather than responding indiscriminately to all surrounding neighbors, animals appear to extract only a small subset of behaviorally relevant information from their social environment. Hidden interaction networks reconstructed from schooling fish reveal that information propagates through highly heterogeneous visual connections Rosenthal et al. [2015], Poel et al. [2021]. Likewise, data-driven models show that coordinated collective behavior can be reproduced when individuals interact with only one or two dynamically selected neighbors Jiang et al. [2017], Lei et al. [2020]. More recently, experimental analyses have further shown that changes in swimming speed dynamically reshape the social interaction network by modulating which neighbors become behaviorally influential, thereby linking locomotor dynamics to the continuous redistribution of social influence within the school Puy et al. [2024]. Together, these studies suggest that collective coordination depends less on the quantity of available social information than on the mechanisms through which that information is filtered, weighted and integrated.

This emerging perspective substantially changes the scientific questions addressed by collective behavior research. The central issue is no longer simply to determine which interaction rules are sufficient to reproduce collective motion, but to identify the computational mechanisms through which animals continuously regulate the information entering these rules. Attraction and alignment should therefore not be regarded as elementary behavioral primitives. Rather, they represent the macroscopic outcome of perceptual and cognitive processes through which individuals evaluate the reliability, relevance and expected consequences of socially acquired information Harpaz et al. [2021], Xiao et al. [2024]. Observable interaction rules become the behavioral signature of hidden computations that remain largely inaccessible through trajectory analysis alone.

This realization exposes one of the principal limitations of current approaches. Despite spectacular progress in behavioral tracking, statistical inference and data-driven modeling, nearly all existing studies remain fundamentally observational. Interaction functions are reconstructed retrospectively from recorded trajectories, while hidden interaction networks are inferred from statistical correlations between neighboring individuals. Such approaches have greatly improved our understanding of collective behavior, yet they cannot uniquely identify the computations generating the observed dynamics. Distinct mechanisms of attention, sensory integration or neighbor selection may produce remarkably similar behavioral trajectories, making causal interpretation inherently difficult. Behavioral observations reveal the consequences of social information processing without directly revealing the algorithms that implement it. Consequently, future progress depends less on collecting increasingly detailed behavioral data than on developing experimental systems capable of manipulating social information itself. Such systems should make it possible to independently manipulate the sensory cues, behavioral signals, and interaction rules exchanged among individuals while preserving the reciprocal feedback that characterizes natural social interactions. Only such experiments can determine which informational variables are actually used by animals to coordinate their behavior.

Recent advances in biomimetic robotics provide an unprecedented opportunity to address this challenge. Artificial fish, birds and insects have progressively evolved from passive moving stimuli into interactive social partners capable of engaging in reciprocal exchanges with living animals Halloy et al. [2007], Faria et al. [2010], De Lellis et al. [2020], Landgraf et al. [2021], Krause et al. [2011], Bonnet et al. [2018], Papaspyros et al. [2023]. Through closed-loop control, robotic agents continuously update their behavior according to the movements of surrounding individuals, allowing stable hybrid groups to emerge. These studies have demonstrated that robots can influence leadership, cohesion, navigation and collective decision-making while offering levels of repeatability and experimental control that cannot be achieved using only living animals Ijspeert et al. [2026]. However, most previous studies have pursued a common objective. Robotic agents were primarily designed to maximize social acceptance by reproducing the appearance and behavior of natural conspecifics as faithfully as possible Butail et al. [2014], Landgraf et al. [2016], Cazenille et al. [2018], Papaspyros et al. [2019]. Their success was therefore evaluated according to their capacity to integrate into animal groups and elicit natural behavioral responses Papaspyros et al. [2024]. Although this biomimetic strategy has considerably advanced animal-robot interaction, it has also tended to obscure what may ultimately be the most important scientific contribution of closed-loop robotics.

We argue that the principal value of robotic partners does not reside in their ability to imitate animals, but in their ability to manipulate social information with a level of precision unattainable in natural systems. Unlike living conspecifics, whose morphology, locomotion, perception and behavior are intrinsically coupled, robotic agents allow these variables to be modified independently. Swimming speed, turning dynamics, responsiveness, neighbor position, behavioral contingency and decision rules can each be controlled experimentally while all remaining aspects of the interaction remain unchanged. Robots therefore become programmable generators of social information rather than merely artificial animals.

This conceptual shift transforms the role of robotics within behavioral biology. Closed-loop robots should not simply be viewed as substitutes for living individuals but as quantitative experimental probes capable of testing explicit hypotheses concerning distributed information processing. Behavioral models cease to be descriptive tools used after experiments have been completed. Instead, they become active components of the experiment itself. Every behavioral algorithm implemented by the robot constitutes a mechanistic hypothesis whose biological validity can be evaluated directly through reciprocal interactions with freely behaving animals. Rather than inferring computational mechanisms solely from collective behavior, closed-loop biohybrid experiments enable candidate algorithms to be implemented directly and their consequences for social interactions to be quantified in real time.

Within this framework, sparse interaction networks acquire a new interpretation. If individuals continuously update the identity of the neighbor exerting the greatest behavioral influence, then collective coordination depends upon an active process of information filtering rather than on static interaction rules Jiang et al. [2017], Lei et al. [2020]. Dynamic social filtering becomes the mechanism through which animals regulate both the amount and the quality of social information entering their behavioral decisions. Such a mechanism naturally reconciles several observations that previously appeared disconnected, including the existence of influential neighbors, the context dependence of interaction strengths, the remarkable efficiency of sparse interaction networks and the coexistence of robust collective organization with rapid behavioral flexibility.

The present study was designed to test this hypothesis directly. We ask whether reciprocal interactions with a closed-loop robotic partner can reveal how living fish dynamically allocate social influence during collective coordination. Using a biomimetic robotic fish whose behavior is continuously updated from the real-time movements of a freely swimming conspecific Papaspyros et al. [2023], we manipulate the informational content of social interactions while preserving their reciprocal nature. This approach enables us to quantify directly how behavioral influence evolves during social interactions and to test whether neighbor selection depends primarily on instantaneous behavioral relevance rather than on spatial proximity alone.

Our results demonstrate that schooling fish continuously reassign social influence among neighboring individuals according to their behavioral relevance, revealing a dynamic social filtering mechanism that provides a causal explanation for the sparse interaction networks inferred previously from observational studies Jiang et al. [2017], Lecheval et al. [2018], Lei et al. [2020], Xue et al. [2023]. Moreover, they show that closed-loop robotic interactions can move beyond reproducing natural behavior to become quantitative experimental tools for identifying the algorithms through which animal groups acquire, filter and propagate social information.

## 2 Results

The closed-loop biohybrid platform enabled a direct comparison between biological groups, biohybrid groups, and numerical simulations using a common set of behavioral observables (Fig. 1A and SI Appendix Fig. S1). Individual behavior was quantified by the distance to the wall *r*_w_ and the wall-incidence angle *θ*_w_, which characterize interactions with the physical environment (Fig. 1C). The consequences of social interactions were assessed from the resulting spatial organization and movement coordination of the group using the inter-individual distance *d*, the viewing angle *ψ*, and the relative orientation Δ*ϕ* (Fig. 1D). These complementary observables were subsequently used to evaluate the ability of the robotic fish to reproduce both individual behavior and the collective dynamics emerging from social interactions.

**Figure 1:**
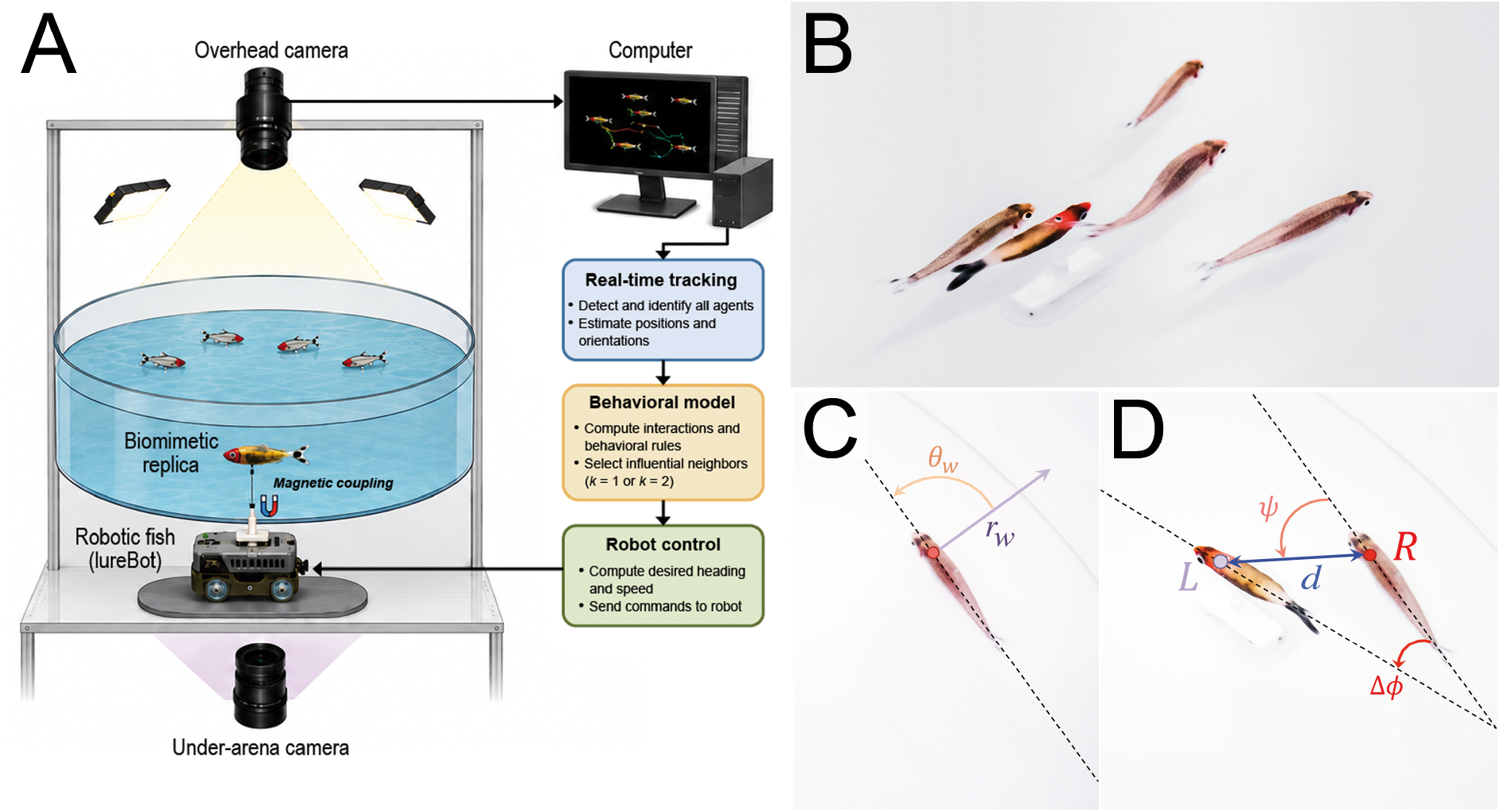
Closed-loop biohybrid platform and behavioral observables used to quantify social interactions. (A) Schematic representation of the closed-loop biohybrid platform used for causal testing of social interaction rules. Freely swimming *Hemigrammus rhodostomus* were continuously tracked in real time while interacting with an autonomous robotic fish carrying a biomimetic replica. Fish positions and orientations were used to update a behavioral controller implementing spontaneous locomotion, wall avoidance, and social interactions. The computed motor commands were transmitted to the robotic fish, whose movements continuously modified the sensory environment of the biological group, establishing reciprocal closed-loop interactions throughout each experiment. (B) Photograph of a school of *H. rhodostomus* swimming with the biomimetic robotic lure during a closed-loop biohybrid experiment. (C) Individual behavioral observables. Individual swimming behavior was quantified by the distance to the arena wall *r*_w_ and the angle of incidence to the wall *θ*_w_, defined as the angle between the swimming direction and the local radial direction. (D) Collective behavioral observables. Pairwise interactions were characterized by the inter-individual distance *d*, the perception angle *ψ*, corresponding to the angular position of a neighbor within the focal fish’s visual field, and the relative orientation Δ*ϕ* between the headings of two individuals (L: LureBot; R: Real Fish). These geometric variables were used both to reconstruct the interaction functions from experimental trajectories and to quantify social coordination in biological, biohybrid, and simulated groups.

### 2.1 Closed-loop robotic fish accurately reproduces the behavior of isolated individuals

The first objective of this study was to determine whether the behavioral model implemented in the robotic fish accurately reproduced the spontaneous locomotion of *Hemigrammus rhodostomus* in the absence of social interactions. This validation was essential because any discrepancy at the individual level would inevitably propagate to the collective dynamics. Both isolated fish and the robotic fish displayed the characteristic burst-and-coast locomotion of the species, consisting of alternating acceleration phases and passive gliding trajectories (Fig. 2C,D; Movie S1). In both cases, trajectories were dominated by persistent circular motion along the arena boundary, reflecting the interaction between intermittent propulsion and wall avoidance rather than attraction towards the wall itself (Fig. 2A,B). The robot reproduced this behavior with remarkable fidelity, generating trajectories visually indistinguishable from those of living fish except for a lower frequency of spontaneous U-turns.

**Figure 2:**
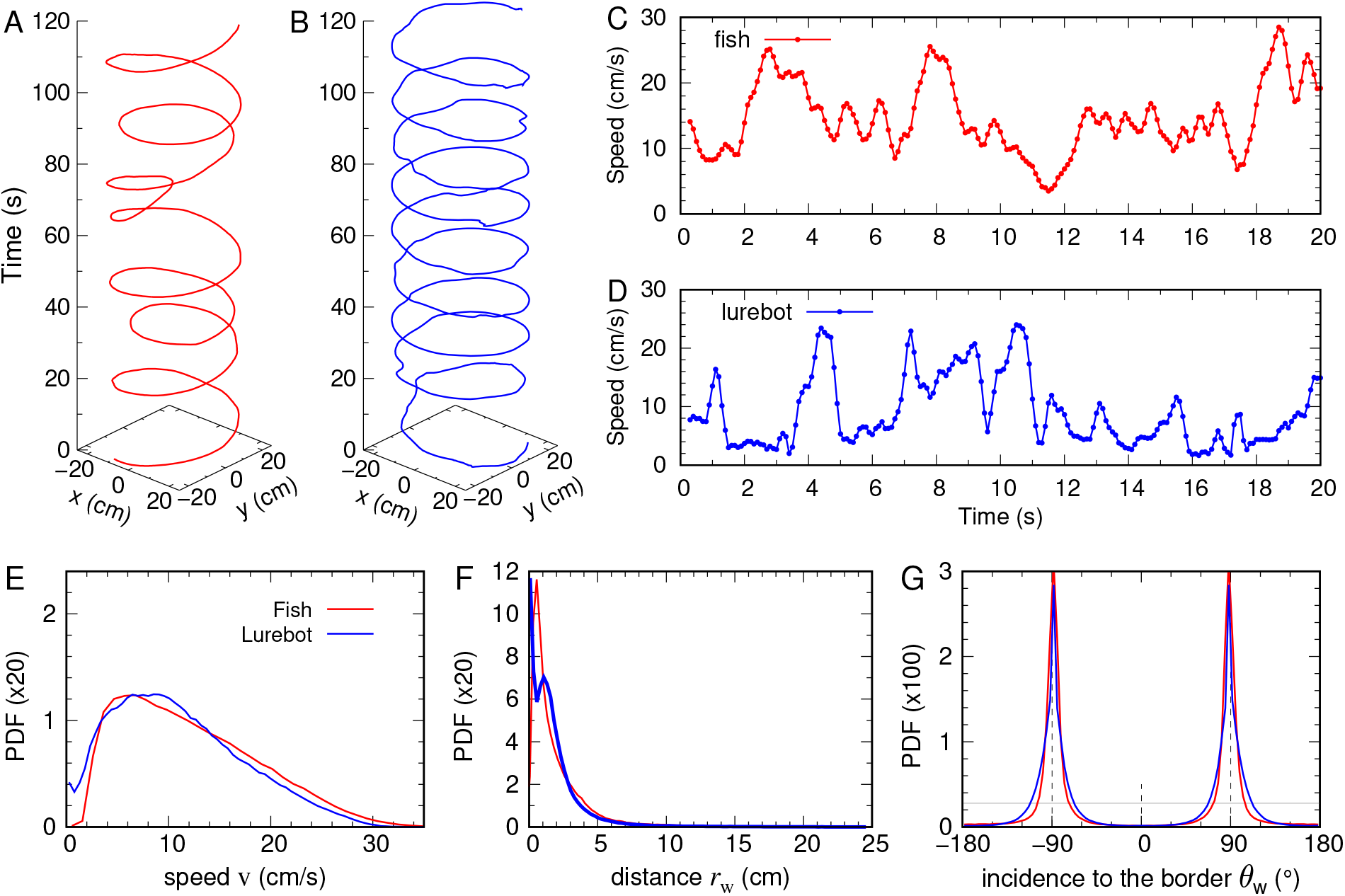
The robotic fish reproduces the spontaneous locomotion of a solitary fish (*N* = 1). (A, B) Representative examples of individual trajectories of fish (red) and LureBot (blue) over 120 s. (C, D) Speed-profiles displaying burst-and-coast dynamics over 20 s. (E) Probability density function (PDF) of speed, (F) distance to the border *r*_w_, and (G) angle of incidence to the border *θ*_w_. Horizontal gray line in (G) denotes uniform PDF.

Quantitative comparisons confirmed this agreement. The probability distributions of swimming speed (Fig. 2E), distance to the wall (Fig. 2F), and incidence angle relative to the arena boundary (Fig. 2G) closely matched those measured experimentally. Mean swimming speed reached 11.0 ± 6.3 cm/s for the robotic fish compared with 12.2 ± 6.8 cm/s for the isolated fish, while both agents maintained nearly identical distances from the arena wall (3.7 ± 3.8 cm for the robotic fish compared to 4.5 ± 3.5 cm for the isolated fish) and followed predominantly tangential trajectories. These results demonstrate that the behavioral controller of the robot correctly reproduces both the locomotor statistics and the interaction with the physical environment that characterize isolated fish. Numerical simulations of the model produced similar spatial statistics (SI Appendix, Fig. S2). Consequently, all subsequent biohybrid experiments relied on parameters specifically calibrated for the physical robot while preserving the structure of the behavioral model.

### 2.2 A robotic fish implementing the interaction model integrates into fish pairs

Having established that the robotic fish accurately reproduces individual behavior, we next investigated whether the same behavioral controller generated natural social interactions when facing a freely swimming conspecific.

Biohybrid pairs rapidly formed cohesive groups whose trajectories closely resembled those observed in pairs of living fish (SI Appendix, Fig. S3 and Movie S2). Throughout the experiments, the robotic fish continuously adjusted its movements according to the position and orientation of its biological partner, while the fish reciprocally modified its own trajectory in response to the robot. This reciprocal closed-loop interaction generated stable coordinated swimming over the entire duration of the trials, demonstrating that the behavioral controller supports genuine social interactions rather than simple stimulus following.

The behavioral similarity between both agents was remarkably high (Fig. 3). The swimming-speed distributions of the robot and the biological fish were nearly indistinguishable, indicating that the fish continuously adapted its own locomotion to match that of the robotic partner (Fig. 3A). Both individuals converged toward the same mean swimming speed (10.6 ± 5.7 cm/s), providing direct evidence of mutual behavioral adjustment. This common swimming speed was slightly lower than that measured in control pairs composed of two biological fish, suggesting that the interaction with the robot shifted the collective dynamics toward a slower but highly coordinated swimming regime. Likewise, both individuals explored the arena using nearly identical spatial strategies, remaining at comparable distances from the wall (4.4 ± 3.8 cm) and exhibiting similar wall-incidence angle distributions (Fig. 3B,C). These observations demonstrate that the robotic fish preserves the natural balance between locomotion, wall avoidance, and social interactions that characterizes freely swimming fish.

**Figure 3:**
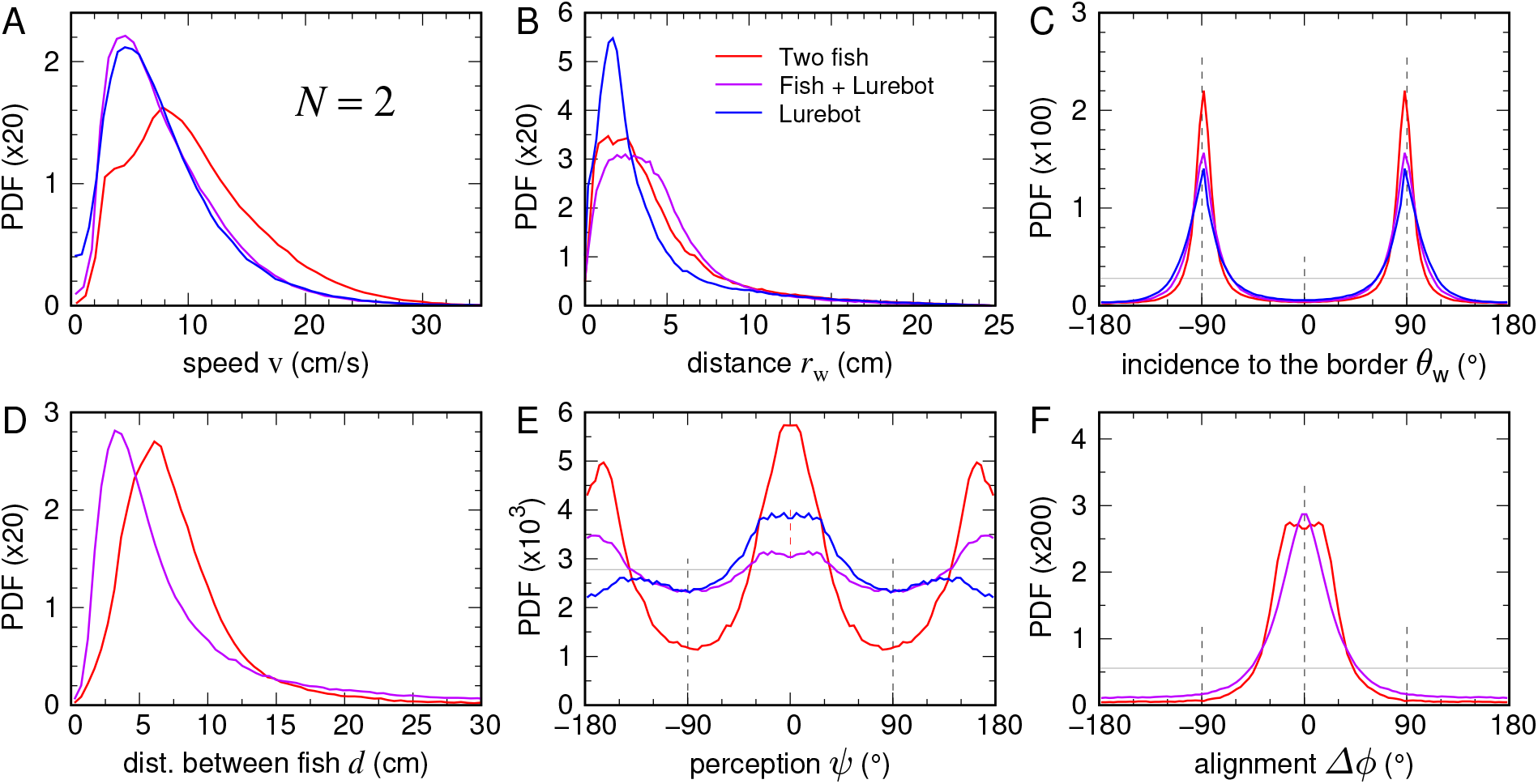
Closed-loop robotic fish reproduces natural pairwise coordination (*N* = 2). Probability density function (PDF) of (A) speed v, (B) distance to the border *r*_w_, (C) angle of incidence to the border *θ*_w_, (D) distance between individuals *d*, (E) angle of perception *ψ*, and (G) alignment Δ*ϕ* in pairs of fish (red), compared to biohybrid pairs, fish (magenta) and LureBot (blue). Horizontal gray lines in (C, E, F) denote uniform PDF.

Numerical simulations further supported these observations (SI Appendix, Fig. S3). The behavioral model quantitatively reproduced the experimental distributions of swimming speed, distance to the arena wall, and wall-incidence angle observed in biological fish pairs. At the collective level, simulations also accurately captured the inter-individual distance, the relative orientation between fish, and their spatial organization. This close agreement confirms that the interaction functions reconstructed from trajectory data provide a quantitatively predictive description of pairwise coordination in freely swimming fish.

Collective descriptors further confirmed the successful social integration of the robotic fish while revealing an unexpected modification of the spatial organization of the pair. Relative-orientation distributions remained nearly identical to those measured in biological control pairs, both exhibiting a sharp peak around zero, indicating that the robotic fish reproduced the alignment dynamics of natural fish pairs with high fidelity (Fig. 3F). In contrast, the relative-position distributions revealed a marked difference in spatial organization. Control pairs predominantly adopted an in-line configuration, with one fish swimming behind the other (Fig. 3E). Such leader–follower formations have previously been reported in zebrafish and interpreted as a stable consequence of the combined effects of social interactions and hydrodynamic coupling Porfiri et al. [2021], Burbano-L. and Porfiri [2021]. By contrast, biohybrid pairs spent most of their time swimming side by side. Although the robotic fish occasionally followed its biological partner, the dominant configuration corresponded to a lateral arrangement in which both individuals swam abreast, as illustrated in Movie S2.

This change in spatial organization also explains the shorter mean inter-individual distance measured in biohybrid pairs compared with biological control pairs (Fig. 3D). This lateral arrangement primarily resulted from the behavior of the robotic fish, whose controller actively maintained a side-by-side position relative to its biological partner (see Movie S2). Unlike a living fish, the robotic fish is insensitive to the hydrodynamic disturbances generated by neighboring swimmers. Consequently, while the interaction model is sufficient to reproduce natural levels of cohesion and alignment, the absence of reciprocal hydrodynamic feedback likely stabilizes side-by-side swimming and reduces the tendency to adopt the in-line formations commonly observed in biological fish pairs.

Taken together, the agreement between numerical simulations and biological fish pairs demonstrates that the reconstructed interaction functions provide a quantitatively predictive description of pairwise coordination. The only systematic discrepancy arises in biohybrid pairs, where the robotic fish stabilizes side-by-side swimming. This result indicates that while behavioral interactions are sufficient to generate natural cohesion and alignment, hydrodynamic interactions contribute to shaping the preferred spatial organization of pairs by promoting the leader–follower configurations observed in freely swimming fish.

### 2.3 Closed-loop experiments reveal dynamic social filtering and the complementary role of hydrodynamic interactions

The central prediction of the interaction model is that fish continuously select only a very small subset of neighbors when adjusting their swimming direction. Rather than integrating information from all surrounding individuals, directional decisions are assumed to depend exclusively on the one or two neighbors exerting the strongest instantaneous influence Lei et al. [2020], Xue et al. [2023], Lin et al. [2025]. To test this prediction experimentally, we introduced the robotic fish into groups containing four freely swimming fish and compared two alternative behavioral controllers. In the first controller, the robot interacted exclusively with its single most influential neighbor (*k* = 1). In the second, it simultaneously integrated the contributions of the two most influential neighbors (*k* = 2). These biohybrid experiments provided a direct causal test of the dynamic neighbor-selection hypothesis that could not be achieved from trajectory analyses alone.

Under both interaction hypotheses, the robotic fish rapidly integrated into the biological groups and participated in the spontaneous formation of cohesive schools whose trajectories closely resembled those of groups composed exclusively of living fish (SI Appendix, Fig. S4 and Movie S3). The robot remained embedded within the moving group throughout the experiments rather than occupying peripheral positions or behaving as an external stimulus. Frequent exchanges of neighbors and continuous modifications of the robot’s social partner reflected the highly dynamic interaction network characteristic of schooling fish. These observations demonstrate that the behavioral controller continuously adapts to the evolving social context rather than simply following one particular individual.

Quantitative analyses confirmed these qualitative observations (Figs. 4 and 5). The distributions, as well as the mean values, of swimming speed, distance to the arena wall, and wall-incidence angle remained remarkably similar between biological and biohybrid groups for both interaction hypotheses. Mean swimming speed reached 13.6 ± 7.2 cm/s for *k* = 1 and 15.1 ± 7.4 cm/s for *k* = 2, compared with 15.2 ± 6.0 cm/s in control groups of five biological fish. Likewise, the average distance to the arena wall was 9.9 ± 5.2 cm for *k* = 1 and 9.3 ± 5.0 cm for *k* = 2, remaining close to the value measured in biological groups (7.0 ± 4.0 cm). These results indicate that replacing one fish by the robotic fish does not alter the individual locomotor behavior of the remaining animals, independently of the amount of social information processed by the robot.

**Figure 4:**
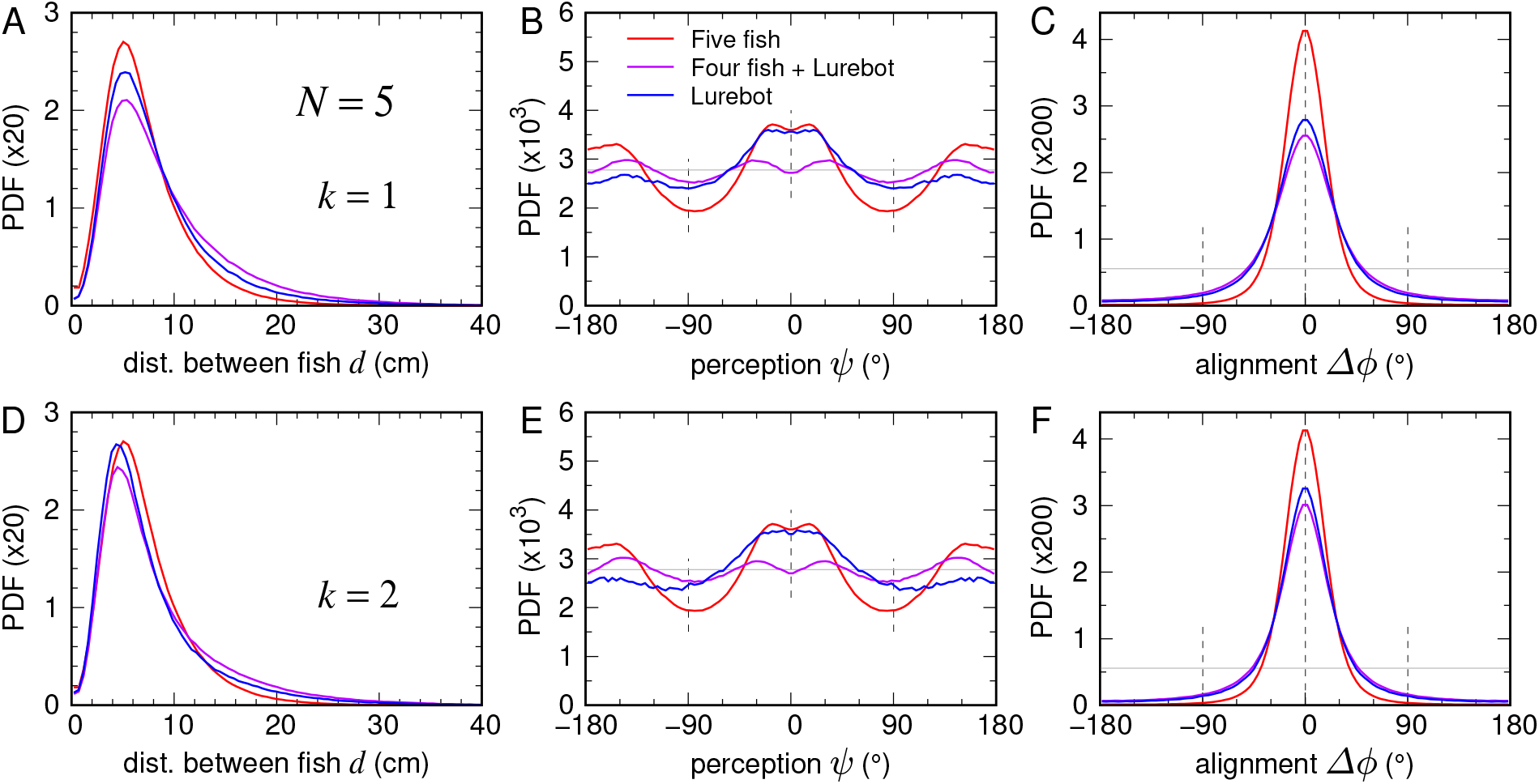
Dynamic neighbor selection accurately reproduces pairwise interaction geometry (*N* = 5). Probability density function (PDF) of (A, D) distance between individuals *d*, (B, E) angle of perception *ψ*, and (C, F) alignment Δ*ϕ* in groups of five fish (red), compared to biohybrid groups of four fish (magenta) and a LureBot (blue), when the LureBot interacts with its (A–C) *k* = 1 or (D–F) *k* = 2 most influential neighbors. Horizontal gray lines in (B, C, E, F) denote uniform PDF.

**Figure 5:**
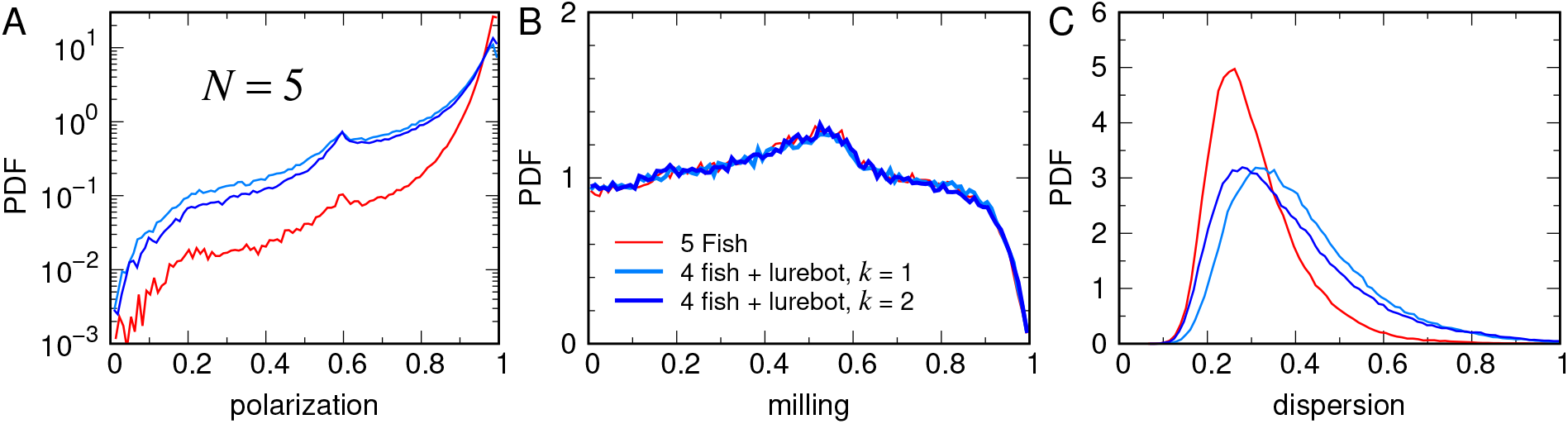
A single influential neighbor is sufficient to reproduce collective organization. Probability density function (PDF) of (A) polarization, (B) milling, and (C) dispersion in groups of five fish (red), compared to biohybrid groups of four fish and a LureBot interacting with its *k* = 1 (light blue) or *k* = 2 (dark-blue) most influential neighbors.

Collective descriptors likewise showed a high level of agreement between biological and biohybrid groups. Nearest-neighbor distance remained close to the biological value for both interaction hypotheses, reaching 9.1 ± 5.9 cm for *k* = 1 and 8.5 ± 6.0 cm for *k* = 2, compared with 7.1 ± 4.0 cm in control groups. Alignment also exhibited only a slight increase when the robot interacted with two neighbors instead of one (Fig. 4). By contrast, the spatial organization of the schools remained essentially unchanged. The distributions of viewing angles were nearly identical for *k* = 1 and *k* = 2, indicating that the number of influential neighbors had virtually no effect on the relative positioning of individuals within the group. Compared with biological groups, however, biohybrid schools displayed a greater tendency to adopt side-by-side configurations, whereas biological fish more frequently formed weak leader–follower arrangements. Although this preference was substantially less pronounced than in isolated fish pairs, it was consistently observed across all biohybrid experiments. Movie S3 indicates that this configuration primarily results from the robotic fish, which preferentially maintains a lateral position relative to nearby fish while occasionally swimming behind one of them.

Global collective properties exhibited the same robustness. Mean milling remained very low and was identical in biological and biohybrid groups (⟨*M*⟩ = 0.47), indicating that none of the experimental conditions produced coherent rotational motion (Fig. 5B). Mean polarization was only slightly higher in biological schools (⟨*P*⟩ = 0.96) than in biohybrid groups (⟨*P*⟩ = 0.86 for *k* = 1 and ⟨*P*⟩ = 0.88 for *k* = 2), while remaining consistently close to the fully polarized state (Fig. 5A). Group dispersion was only marginally higher for *k* = 1 (0.4 ± 0.1) than for *k* = 2 (0.4 ± 0.2), whereas biological groups exhibited a value of 0.3 ± 0.1 (Fig. 5C). Altogether, these observations show that restricting the robot to a single influential neighbor is already sufficient to reproduce the large-scale organization and coordination of natural fish schools. Incorporating a second influential neighbor produces only modest quantitative improvements without qualitatively modifying the collective dynamics.

Comparison with numerical simulations further clarified the origin of these small differences (SI Appendix, Fig. S5). The simulations using *k* = 2 reproduced the experimental observations more accurately than the simulations using *k* = 1. In particular, nearest-neighbor distance was accurately reproduced under both interaction hypotheses, with mean values of 7.3 ± 4.9 cm for *k* = 1 and 7.4 ± 4.8 cm for *k* = 2, closely matching the value measured in biological schools (7.1 ± 4.0 cm). Likewise, the distributions of viewing angle and relative orientation, together with group dispersion (0.3 ± 0.1), agreed more closely with the experimental measurements when two influential neighbors were considered. These results suggest that hydrodynamic interactions, whose effects are implicitly incorporated into the interaction functions reconstructed from biological trajectories but do not influence the robotic fish itself, contribute to the subtle quantitative differences observed between the two interaction hypotheses. Consequently, reproducing the exact cohesion and polarization of biological schools in numerical simulations requires each fish to interact with its two most influential neighbors, whereas the robotic fish achieves comparable collective coordination by responding only to its single most influential neighbor.

Taken together, these experiments provide direct causal evidence that collective coordination in schooling fish does not emerge from integrating increasing amounts of social information. Instead, it relies on a dynamic filtering mechanism through which behavioral decisions are continuously based on the currently two most influential neighbors. Because the identity of these neighbors changes rapidly as the group reorganizes, each fish successively interacts with many different group members while processing only two social interactions at a time. This remarkably simple decision strategy is sufficient to reproduce the collective dynamics of natural fish schools across multiple levels of biological organization.

## 3 Discussion

Understanding how individuals process social information to generate coordinated collective behavior remains a central challenge in behavioral biology Camazine et al. [2001], Sumpter [2010], Couzin [2009]. The present study provides direct experimental evidence that the interaction rules reconstructed from behavioral trajectories capture the essential mechanisms governing collective coordination in schooling fish. By combining a closed-loop robotic fish with freely swimming conspecifics, we moved beyond the traditional validation of behavioral models based solely on statistical agreement between numerical simulations and experiments. Instead, the inferred interaction rules were implemented in an autonomous robotic partner whose behavior continuously influenced, and was influenced by, living fish under natural social conditions. Although quantitative differences remained between numerical simulations and biohybrid experiments, particularly in the spatial organization and cohesion of groups, all three approaches consistently reproduced the principal behavioral signatures observed from isolated individuals to pairs and small schools. These complementary results demonstrate that the proposed interaction framework possesses substantial predictive power while simultaneously revealing the respective contributions of behavioral and physical interactions to collective organization. Most importantly, our experiments provide the first causal validation that dynamic social filtering, whereby behavioral decisions are continuously based on the currently most influential neighbor, constitutes a sufficient mechanism to generate coordinated collective motion in schooling fish.

The present findings substantially strengthen the emerging view that collective coordination relies on selective rather than exhaustive processing of social information. Earlier studies reconstructed interaction networks from behavioral trajectories and showed that only a limited number of neighbors contribute significantly to the behavioral decisions of each individual Rosenthal et al. [2015]. Likewise, computational analyses predicted that coordinated schooling behavior could be reproduced if each fish interacted only with the one or two neighbors exerting the strongest instantaneous influence Jiang et al. [2017], Lei et al. [2020]. More recently, dynamic social interactions have been shown to depend strongly on swimming speed, suggesting that influential neighbors emerge from continuously changing behavioral contexts rather than from persistent leadership relationships Puy et al. [2024], Escobedo et al. [2026]. Our biohybrid experiments extend these observations by providing direct causal evidence that processing the behavior of a single or two dynamically selected neighbors is sufficient to reproduce the organization of natural fish schools. Importantly, interacting with a single or two neighbors does not imply following the same individuals over extended periods. On the contrary, the identity of the most influential neighbor changes continuously as the group reorganizes, allowing the robotic fish to successively exchange information with many different group members while processing only one or two social interactions at a time. This dynamic filtering mechanism therefore reconciles two apparently contradictory properties of animal groups. It dramatically reduces the amount of information that must be processed by each individual while preserving the flexibility and robustness required for coherent collective behavior in rapidly changing social environments.

The only systematic difference between biological and biohybrid groups concerned their relative spatial organization, providing new insight into the interplay between social and hydrodynamic interactions during collective swimming. In both pairs and groups of five fish, replacing one individual by the robotic fish increased the occurrence of side-by-side swimming while preserving natural levels of cohesion, alignment and polarization. At first sight, this result appears surprising because previous experimental and theoretical studies on zebrafish concluded that in-line swimming represents the most stable configuration for burst-and-coast swimmers. Small perturbations away from this leader–follower arrangement are dynamically compensated through the combined action of social interactions and hydrodynamic coupling, causing fish to return spontaneously to an in-line formation Porfiri et al. [2021]. Likewise, models incorporating multiple sensory pathways have shown that locomotor decisions emerge from the integration of visual and hydromechanical information, both of which contribute to the stability of collective swimming Burbano-L. and Porfiri [2021]. This interpretation is further supported by the recent RoboTwin platform, which experimentally demonstrated that robotic replicas reproducing the body kinematics of real fish can be used to isolate and quantify the contribution of hydrodynamic interactions to collective swimming Li et al. [2024]. Our biohybrid experiments provide a simple explanation for why biohybrid groups preferentially adopt side-by-side swimming, despite previous theoretical and experimental studies showing that in-line swimming is the most stable configuration for burst-and-coast fish. Unlike a living fish, the robotic fish continuously responds to social interactions but remains insensitive to the hydrodynamic disturbances generated by neighboring fish, whose influence becomes most pronounced when individuals swim in close proximity. Moreover, the robotic platform has a limited ability to reproduce the fine-scale adjustments continuously performed by living fish. The magnetic coupling between the robot and the biomimetic lure, together with the mechanical inertia of the system and the open-loop execution of each burst once initiated, likely contributes to the remaining differences between biohybrid and biological groups. Consequently, the robot actively maintains a lateral position relative to nearby fish, whereas the fluid-mediated feedback that naturally favors in-line configurations is absent. This difference explains why side-by-side swimming becomes more frequent in biohybrid groups without affecting the overall coordination of the school. The fact that this spatial organization is observed both in pairs and in groups of five individuals further indicates that it primarily results from the behavior of the robotic fish rather than from a modification of the interaction strategy adopted by the living fish.

The comparison between biohybrid experiments and numerical simulations further clarifies the respective contributions of behavioral and hydrodynamic interactions. Together, these complementary approaches highlight how behavioral and hydrodynamic interactions jointly contribute to the organization of fish schools. The behavioral interaction functions used throughout this study were reconstructed from trajectories of freely swimming fish and therefore implicitly incorporate the behavioral consequences of the hydrodynamic interactions experienced by real animals. However, one important limitation of this approach is that fish rarely swim at very short inter-individual distances, resulting in relatively few observations below approximately 3 cm from which interaction functions can be estimated Calovi et al. [2018]. Consequently, the reconstructed interaction functions are likely less accurate in this regime and probably underestimate the contribution of hydrodynamic interactions when neighboring fish swim in close proximity. A complete mechanistic description would ultimately require coupling the reconstructed behavioral interaction rules with an explicit model of the surrounding flow together with a description of how fish sense and respond to hydrodynamic cues. Neither the numerical simulations nor the robotic controller incorporate such an explicit representation of hydrodynamic interactions, relying instead on behavioral interaction functions reconstructed from experimental trajectories. Despite this limitation, the behavioral model reproduces remarkably well the collective dynamics and spatial organization of biological schools. Accordingly, when all individuals in the numerical simulations are governed exclusively by this behavioral model, considering the two most influential neighbors produces the closest quantitative agreement with the experimental observations, reproducing more accurately the cohesion, polarization and neighbor distributions measured in biological schools, consistent with previous computational analyses Jiang et al. [2017], Lei et al. [2020], Xue et al. [2023], Lin et al. [2025]. In the biohybrid experiments, however, only the robotic fish does not physically experience the fluid-mediated interactions generated by neighboring swimmers, whereas the four biological fish continue to experience both behavioral and fluid-mediated interactions. Under these conditions, allowing the robot to respond exclusively to its single most influential neighbor is sufficient to maintain collective cohesion and polarization at levels nearly identical to those observed in entirely biological groups. Because the robotic fish represents only one of the five individuals in the group, its interaction strategy affects only a limited fraction of the collective interaction network. By contrast, in the numerical simulations the same interaction rule is applied to every individual, making the choice of *k* much more influential for the collective dynamics. These results therefore indicate that the robotic fish can simplify its behavioral decisions to a single influential neighbor without measurably altering the collective organization of the group.

More generally, these findings emphasize that collective swimming emerges from the continuous interaction between behavioral decisions and the physical environment. They also illustrate the unique potential of closed-loop biohybrid experiments for disentangling mechanisms that remain intrinsically coupled in natural animal groups. By selectively removing one component of the interaction process from a single individual while preserving reciprocal interactions with the rest of the group, this approach provides a powerful experimental framework for quantifying the respective roles of social behavior and physical coupling in the emergence of collective organization.

Our study also demonstrates that the proposed interaction framework remains predictive across multiple levels of biological organization. Behavioral models calibrated from pairwise interactions often reproduce dyadic behavior but rapidly lose predictive power when extrapolated to larger groups, where increasingly complex interaction networks may give rise to emergent collective phenomena Vicsek and Zafeiris [2012], Herbert-Read [2016]. In contrast, the same behavioral rules accurately reproduced spontaneous locomotion and wall-following behavior in isolated fish, social coordination in pairs, and the organization of groups containing five individuals without introducing any additional behavioral mechanisms. The close agreement observed between numerical simulations, biohybrid experiments and entirely biological groups therefore demonstrates that the proposed interaction model possesses genuine predictive power rather than merely descriptive accuracy. More importantly, the successful integration of the robotic fish cannot be explained solely by statistical similarities between simulated and experimental trajectories. Throughout the experiments, the robot continuously exchanged information with living fish under reciprocal closed-loop interactions, demonstrating that both agents responded to compatible behavioral cues in real time. These results support the view that the mechanisms governing attraction, alignment, spontaneous fluctuations and interactions with the physical environment constitute a unified behavioral framework capable of explaining collective coordination across several levels of biological organization.

Beyond the biological conclusions, our work also illustrates the methodological potential of closed-loop biohybrid robotics for studying collective behavior. Most previous studies integrating robots into fish schools primarily sought to maximize the social acceptance of artificial conspecifics by improving their morphology, appearance or locomotor realism, or by implementing increasingly sophisticated behavioral controllers Krause et al. [2011], Bonnet et al. [2018], Landgraf et al. [2021, 2016]. Likewise, model-based robotic control has previously been proposed as a powerful strategy for testing behavioral hypotheses under closed-loop conditions De Lellis et al. [2020]. Our study extends this approach by shifting the objective from social integration itself to the experimental validation of behavioral mechanisms. Rather than asking whether a robotic fish can be accepted as a conspecific, we use the robot as a programmable experimental agent whose interaction rules can be manipulated independently of morphology, sensory cues and environmental conditions. This transformation of the robot from a biomimetic stimulus into a quantitative experimental instrument provides a direct causal framework for testing behavioral models under fully reciprocal interactions. More generally, the same strategy could readily be extended to investigate how collective decisions emerge when only a subset of individuals possesses information, when interaction rules change dynamically, or when different sensory modalities contribute simultaneously to collective coordination.

Our findings have broader implications for understanding collective intelligence in biological systems Camazine et al. [2001], Sumpter [2010], Couzin [2009], McMillen and Levin [2024], Moussaïd et al. [2009]. Animal groups are often viewed as distributed information-processing systems whose performance increases as more individuals contribute information. The present results suggest a different perspective. Efficient collective coordination does not require each individual to integrate an ever-growing amount of social information. Instead, behavioral decisions appear to rely on a dynamic filtering process that continuously identifies the socially most relevant neighbor while rapidly updating this choice as the interaction network evolves. Such a strategy dramatically reduces the computational demands imposed on individuals while preserving the flexibility, responsiveness and robustness required for coherent collective behavior. Dynamic social filtering may therefore represent a general computational principle through which decentralized biological systems reconcile cognitive simplicity with collective efficiency. Because the experimental framework introduced here allows behavioral rules to be manipulated independently under reciprocal closed-loop interactions, it opens new opportunities for investigating whether similar mechanisms underlie information processing in other animal societies, including bird flocks, insect colonies and mammalian groups, as well as in bio-inspired robotic swarms.

## Materials and Methods

### Ethics statement

Experiments were approved by the Ethical Committee for Animal Experimentation of the Toulouse Biology Research Federation (C2EA-01) and were performed in an approved fish facility (A3155501) under permit APAFIS#27303-2020090219529069 v8 in agreement with the French legislation. All procedures were designed to minimize stress and handling. Fish were transferred from rearing tanks to the experimental setup with minimal manipulation. Each individual was used in only one one-hour experimental session per day. Swimming ability was monitored throughout; fish exhibiting impaired or absent swimming activity were excluded and replaced. No animals were sacrificed during this study.

### Study species

Experiments were performed on the rummy-nose tetra (*Hemigrammus rhodostomus*), a highly social freshwater fish widely used as a model system for studying collective motion. Adult fish (body length 35 ± 3 mm) were obtained from a commercial supplier (Amazonie Labège, Toulouse, France) and maintained in groups in 16 L aquariums on a 12:12 hour, dark:light photoperiod, at 27.0 ± 0.8^◦^ C and were fed *ad libitum* with fish flakes.

### Experimental setup

Experiments were performed using a biomimetic robotic platform previously developed for real-time interactions with freely swimming fish Papaspyros et al. [2023]. The platform consists of an autonomous mobile robot (LureBot) carrying a three-dimensional biomimetic replica of *H. rhodostomus* that reproduces the external appearance of a conspecific (Fig. 1A and SI Appendix, Fig. S1). The robot moved beneath the experimental arena while the replica was magnetically coupled above the arena floor, allowing unrestricted planar motion without disturbing the fish.

The position of every fish was continuously tracked using an overhead monochrome camera (Basler acA4024-29uc, 4024 × 3036 pixels, 30 Hz; downsampled to 640 × 480 for the tracking algorithm). A second camera positioned beneath the arena simultaneously tracked the robot’s position and orientation using two differently colored LEDs mounted at the front and rear of the LureBot. Real-time image processing combined information from both cameras to identify all biological and artificial agents and continuously update the behavioral controller implemented in the robot. This closed-loop architecture allowed reciprocal interactions between the robotic fish and freely swimming individuals throughout the experiments.

### Experimental procedure

Experiments were conducted in a circular arena (radius 25 cm) filled with approximately 5 cm of water obtained directly from the maintenance aquarium to preserve identical physicochemical conditions. Water temperature was maintained at 27^◦^ C throughout all experiments.

Fish were randomly selected from one of the housing tanks and introduced into the arena, where they acclimated for 15 min before recordings began. During acclimation, the robotic fish remained stationary. Individuals exhibiting abnormal stress-related behavior were replaced before the beginning of the experiment. Each trial lasted one hour. After testing, fish were transferred to a temporary holding tank to prevent repeated participation on the same day. Fish were randomly sampled from the housing tanks and tested on different days, ensuring that no individual participated in experiments on consecutive days and thereby minimizing habituation to the robotic fish.

Three experimental conditions were investigated. First, isolated fish and isolated robotic fish were recorded to characterize spontaneous locomotion and interactions with the arena boundary. Second, pairwise interactions were examined by comparing control groups composed of two living fish with biohybrid groups composed of one living fish and one robotic fish. Third, collective interactions were investigated in groups containing five individuals. Control groups consisted of five living fish, whereas biohybrid groups contained four living fish and one robotic fish. Two behavioral controllers were tested. In the first condition, the robotic fish interacted exclusively with its most influential neighbor (*k* = 1). In the second condition, behavioral decisions incorporated the two most influential neighbors (*k* = 2). Experimental durations were 10 h for the biological control groups and the biohybrid pairs (*N* = 2), and 8 h for each biohybrid group condition (*N* = 5) with the robotic fish interacting with either its most influential neighbor (*k* = 1) or its two most influential neighbors (*k* = 2).

#### Behavioral tracking and trajectory processing

Videos were analyzed using idtracker.ai, version 4 Romero-Ferrero et al. [2019], providing automatic identity tracking of all individuals with a reported accuracy exceeding 99.5%. Remaining identity swaps and missing detections were corrected using a dedicated post-processing pipeline. Only periods during which all agents were actively swimming were retained for analysis. Frames corresponding to swimming speeds below one body length per second were discarded to remove inactive periods and rare losses of magnetic coupling between the robot and its biomimetic replica. Trajectories were subsequently resampled at 10 Hz (Δ*t* = 0.1 s), reducing tracking noise while preserving the temporal resolution required to quantify social interactions.

#### Quantification of individual and collective behavior

Behavior was quantified using a common set of descriptors characterizing locomotion at the individual, pairwise, and collective levels. The same observables were computed from experimental trajectories, biohybrid experiments, and numerical simulations, allowing direct quantitative comparisons across all conditions.

Individual behavior was characterized by three variables. Swimming speed v was calculated from successive positions of each individual. The distance to the arena wall *r*_w_ was defined as the shortest distance between the fish and the circular boundary of the arena. Wall orientation was quantified by the wall-incidence angle *θ*_w_, corresponding to the angle between the instantaneous swimming direction and the local radial direction. Together, these variables characterize spontaneous locomotion together with interactions with the physical environment (Fig. 1C).

Pairwise social interactions were quantified using three geometric descriptors computed for every pair of individuals (Fig. 1D). The inter-individual distance *d* measured local cohesion. The viewing angle *ψ* corresponded to the angular position of a neighbor within the visual field of the focal fish, whereas the relative orientation Δ*ϕ* quantified the angular difference between the swimming directions of both individuals. These variables constitute the fundamental descriptors from which the social interaction functions implemented in the behavioral model were reconstructed and were used here to quantify social coordination in biological, biohybrid, and simulated groups.

Collective organization in groups of five individuals was quantified using three complementary observables describing directional order, rotational organization, and spatial cohesion:

1. Polarization:

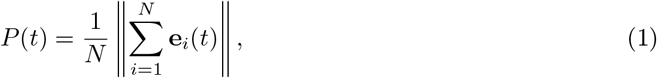

where **e**_*i*_ = **v**_*i*_*/*∥**v**_*i*_∥ is the unit vector in the swimming direction of fish *i*. Polarization measures the degree of alignment within the group and ranges from 0 to 1, with 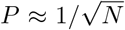 corresponding to disordered motion and *P* = 1 to the fully ordered state where all individuals swim in the same direction.
2. Milling, quantifying the collective rotational motion:

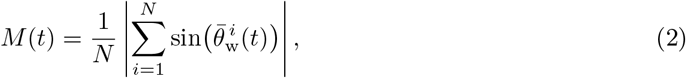

where 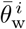 denotes the angle between the swimming direction of fish *i* and its radial position relative to the group centroid. Milling also ranges from 0 to 1, with larger values indicating coherent rotation of the group around its centroid, independently of the direction of rotation.
3. Group cohesion, quantified by the normalized spatial dispersion:

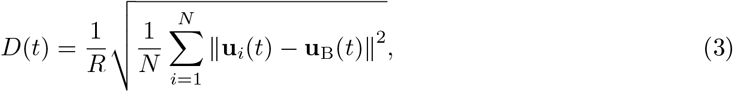

where **u**_*i*_ is the position of fish *i*, **u**_B_ is the group centroid, and *R* is the radius of the experimental arena. Unlike polarization and milling, dispersion is not an order parameter but quantifies the spatial extent of the school, smaller values corresponding to more cohesive groups.

For pairwise experiments, probability density functions (PDFs) were computed for all individual and pairwise descriptors. For groups of five fish, PDFs and corresponding mean values were calculated for all individual, pairwise, and collective observables. Mean values are reported throughout the manuscript as mean ± standard deviation.

#### Behavioral model

The robotic fish was controlled by a quantitative behavioral model previously developed to describe the burst-and-coast locomotion and social interactions of the rummy-nose tetra (*H. rhodostomus*) Calovi et al. [2018], Lei et al. [2020], Xue et al. [2023]. The model was reconstructed from experimental trajectory data and subsequently validated over a broad range of environmental conditions and group sizes. In particular, it accurately reproduces the swimming dynamics of isolated fish as well as the collective organization of groups ranging from pairs to schools of 25 individuals under different illumination levels Xue et al. [2023]. In the present study, the same interaction framework was implemented in the robotic fish to generate real-time closed-loop interactions with freely swimming conspecifics.

The model reflects the burst-and-coast swimming mode characteristic of *H. rhodostomus*. Fish trajectories consist of successive gliding phases separated by short propulsive events, referred to as *kicks*, during which both swimming speed and heading direction are updated. At the *n*-th kick performed by fish *i*, its state is defined by its position 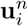 and heading angle 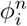. During the kick, the fish selects a new heading,

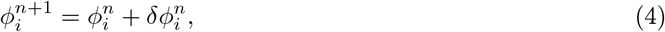

and subsequently glides along a straight trajectory until the next kick occurs. The position at the end of the glide is therefore

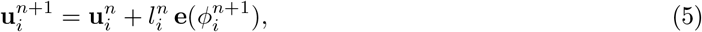

where 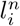 denotes the kick length and **e**(*ϕ*) is the unit vector associated with heading *ϕ*.

Kick duration and initial swimming speed are independently and asynchronously sampled from experimentally measured distributions. Between two consecutive kicks, swimming speed decreases exponentially because of hydrodynamic drag. Consequently, the instantaneous position of the fish during the glide phase is given by

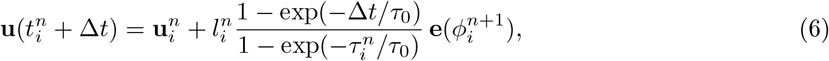

where *τ*_0_ is the characteristic deceleration time and 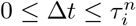.

The directional decision performed at each kick results from the additive contribution of spontaneous behavioral variability, interactions with the physical environment, and social interactions,

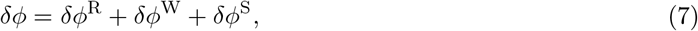

where the superscripts denote the random, wall, and social components, respectively.

The spontaneous component accounts for intrinsic variability in swimming direction and is represented by a Gaussian random process whose amplitude decreases as fish approach the arena boundary,

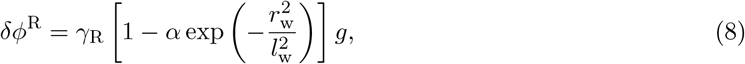

where *r*_w_ is the distance to the wall, *g* is a standard Gaussian random variable, and *γ*_R_, *α*, and *l*_w_ are model parameters.

Interactions with the arena boundary generate a repulsive turning response whose magnitude depends on both the distance to the wall and the fish orientation relative to the boundary,

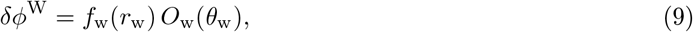

where *f*_w_ describes the decay of wall influence with distance and *O*_w_ captures its angular dependence.

Social interactions combine attraction–repulsion and alignment responses reconstructed directly from experimental trajectory data. The behavioral influence exerted by neighbor *j* on fish *i* depends on their relative distance *d*, viewing angle *ψ*, and relative orientation Δ*ϕ*. The social contribution to heading variation is written as

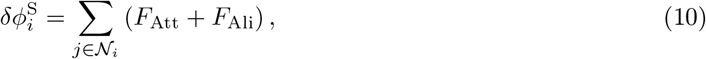

where *F*_Att_ and *F*_Ali_ denote the attraction–repulsion and alignment interactions, respectively. Each interaction is expressed as the product of a distance-dependent function and two angular modulation functions describing anisotropic perception. The complete analytical expressions of these interaction functions are provided in the SI Appendix.

For each behavioral update, neighbors are ranked according to the magnitude of the directional change they induce on the focal fish *i*,

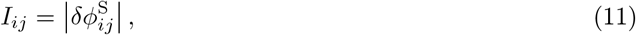

which provides an instantaneous measure of social influence. Depending on the experimental condition, the robotic fish interacted either with the single most influential neighbor (*k* = 1) or with the two most influential neighbors (*k* = 2). This experimental manipulation enabled a direct causal test of the dynamic neighbor-selection hypothesis.

Finally, candidate kicks predicting positions outside the experimental arena were rejected and resampled until a valid trajectory remained inside the circular tank. This rejection procedure preserves the burst-and-coast dynamics while ensuring realistic interactions with the physical boundaries. All parameter values used in the simulations and in the robotic controller are reported in Tables S1-S4.

## Data Availability

The datasets used in the current study are available at: https://doi.org/10.6084/m9.figshare.33086195

## Acknowledgements

This work was partly supported by the Germaine de Staël project no. 2019-17. V. P. and F. M. were also supported by the Swiss National Science Foundation Project ‘Self-Adaptive Mixed Societies of Animals and Robots’, Grant No. 175731. The funders had no role in study design, data collection and analysis, decision to publish, or preparation of the manuscript.

## Supporting Information for

## Burst-and-coast behavioral model

The robotic fish is controlled by a behavioral model previously developed to describe the burst-and-coast motion and social interactions of fish of the species (*Hemigrammus rhodostomus*) Calovi et al. [2018], Lei et al. [2020], Xue et al. [2023]. The interaction functions are reconstructed from experimental trajectory data and have been validated over a broad range of environmental conditions and group sizes, from isolated individuals to schools of 25 fish under different illumination conditions Xue et al. [2023]. In the present study, we use the same interaction model in the robotic fish to generate real-time interactions with freely swimming conspecifics.

### Individual dynamics

The burst-and-coast swimming mode consists in a succession of abrupt impulses (kicks) followed by quasi-passive gliding phases during which the fish decelerates along a nearly straight line.

The state of a fish is determined by its position and velocity vectors **u** = (*x, y*) and **v** = (*v*_*x*_, *v*_*y*_). The heading angle is given by the direction of the velocity, *ϕ* = atan2(*v*_*y*_, *v*_*x*_), and the angle of incidence to the wall by *θ*_w_ = *ϕ* − atan2(*y, x*), so that *θ*_w_ = 0 when the fish swims directly toward the wall. The distance to the wall is thus given by *r*_w_ = *R* − |**u**|, where *R* is the radius of the circular arena.

At the *n*-th kick performed by fish *i*, occurring at time 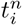, the fish selects a new heading

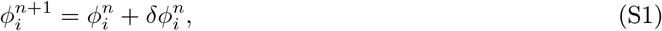

and then glides along a straight segment of length 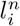. Its position at the end of the glide is given by

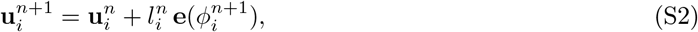

where **e**(*ϕ*) = (cos *ϕ*, sin *ϕ*) is the unit vector in the direction *ϕ*. Kick duration 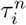 and initial swimming speed are independently and asynchronously sampled from experimentally measured distributions. Between two consecutive kicks, swimming speed decreases exponentially due to hydrodynamic drag, so that the instantaneous position during the glide is

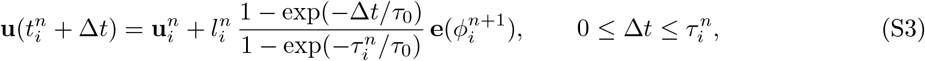

where *τ*_0_ is the characteristic deceleration time.

The heading change *δϕ* performed at each kick is the result of the additive combination of the spontaneous behavioral variability of the fish, the interactions with the physical environment, and the social interactions:

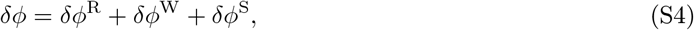

where the superscripts denote the Random, Wall, and Social components, respectively.

The random component *δϕ*^R^ accounts for the intrinsic variability of fish swimming and is represented by a Gaussian random process whose amplitude decreases near the border of the arena:

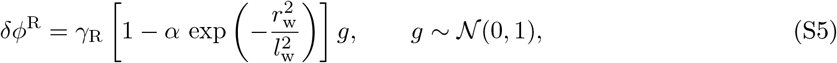

where *l*_w_ is the characteristic range of the influence of the wall, *α* is an attenuation factor, and *γ*_R_ is the amplitude of the noise.

The effect of the wall of the arena *δϕ*^W^ is a repulsive force whose magnitude depends on both the distance to the wall *r*_w_ and the angle of incidence to it, *θ*_w_:

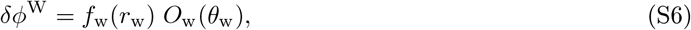

with

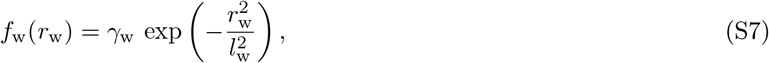

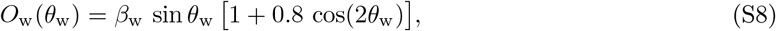

where *γ*_w_ is the intensity and *β*_w_ is a normalization factor for *O*_w_, such that 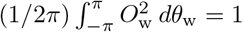.

### Collective dynamics

The social interactions term *δϕ*^S^ combines attraction/repulsion and alignment forces reconstructed directly from experimental trajectory data Calovi et al. [2018]. The behavioral impact of a neighbor *j* of a focal fish *i* depends on their relative state, determined by the distance between them *d*, the viewing angle with which *i* perceives *j, ψ* = atan2(*y*_*j*_ − *y*_*i*_, *x*_*j*_ − *x*_*i*_) − *ϕ*_*i*_, and the relative heading Δ*ϕ* = *ϕ*_*j*_ − *ϕ*_*i*_.

The contribution of the neighbor *j* to the heading variation of *i* is thus given by

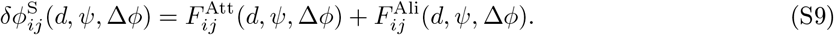

This magnitude quantifies the *influence* exerted by fish *j* on the focal fish *i*:

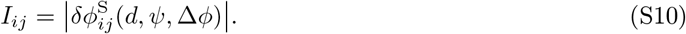

At each kick, all neighbors are ranked according to *I*_*ij*_, and only the *k* most influential ones are retained, typically *k* = 1 or *k* = 2 Lei et al. [2020]. Importantly, the identity of the most influential neighbors can change from one kick to the next, because *I*_*ij*_ depends on the instantaneous relative state of each fish with respect to the focal one.

The final social contribution to heading variation of fish *i* is thus given by:

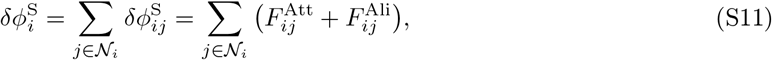

where *N*_*i*_ denotes the set of the *k* most influential neighbors of fish *i*.

The pairwise interactions are factorized in a distance-dependent function and two angular modulation functions. Following Calovi et al. [2018], we distinguish odd (*O*) and even (*E*) angular functions:

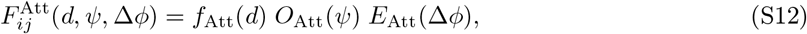

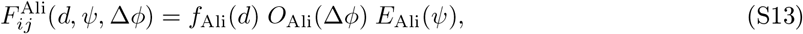

where

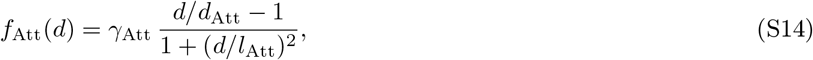

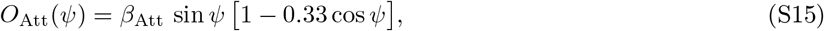

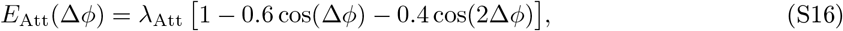

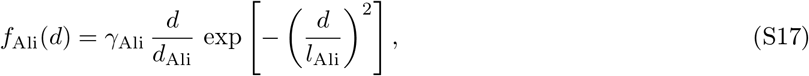

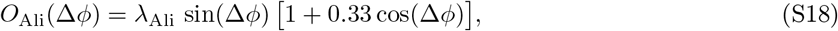

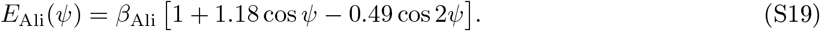

The radial function *f*_Att_ changes from repulsive to attractive when the distance between fish *d* goes from below to above *d*_Att_. The intensity of the attraction is controlled by *γ*_Att_ and the characteristic range of action by *l*_Att_. Similarly, *γ*_Ali_ controls the alignment intensity in the radial function *f*_Ali_, with characteristic length *l*_Ali_. The normalization constants *β*_Att,Ali_ and *λ*_Att,Ali_ are defined analogously to *β*_w_.

### Boundary constraint

When a new kick is calculated, it may happen that the predicted final position of the fish falls outside the circular arena. These tentative kicks are rejected and resampled until a valid one is obtained. This rejection procedure preserves the burst-and-coast dynamics while ensuring realistic boundary interactions.

### Robotic implementation

The same behavioral model is implemented in the LureBot controller to drive real-time, closed-loop interactions with real fish. During the burst phase, the robot computes its target position and accelerates rapidly toward it; during the coast phase, it maintains straight-line motion following the speed profile determined at the onset of its kick.

Unlike the computational model, the robotic implementation must account for physical constraints including physical collision with the border, real-time computation latency and electronic noise. To compensate for these effects, the three parameters *γ*_R_, *γ*_Att_, and *γ*_Ali_ were adjusted independently for the robot and for the numerical simulations. All parameter values are reported in Tables S1-S4.

## Supplementary Note: relation between the robot controller and the simulation model

### Model equations used by the robot controller

At its *n*th kick, the heading of the virtual reference agent is updated according to Eqs. (S1) and (S4). The radial attraction function *f*_Att_ and the odd angular function *O*_Att_ are those of the model, while the remaining interaction functions take the form:

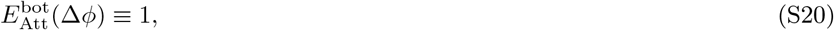

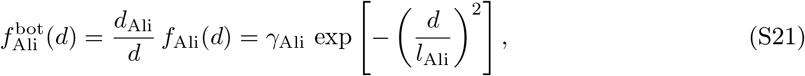

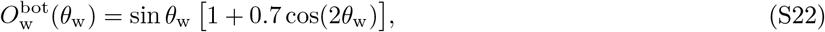

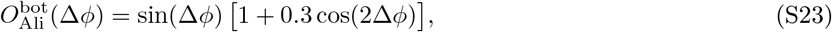

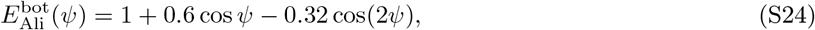

with *β*_w_ = *β*_Att_ = *β*_Ali_ = *λ*_Ali_ = 1, the robot leaving the angular functions un-normalized.

The even function *E*_Att_ is replaced in the robot by its average over the relative heading, ⟨*E*_Att_⟩_Δ*ϕ*_ ≡ 1, so that the attraction exerted by a neighbor depends only on its distance *d* and viewing angle *ψ*. Headings are estimated less accurately than positions by the tracking system, and Δ*ϕ*, being a difference of two such estimates, is the noisiest input available to the controller while the robot is moving; retaining *E*_Att_ would propagate this noise multiplicatively into the attraction, which is the dominant social term. Averaging it leaves the mean attraction strength unchanged, and the dependence of the social response on the relative heading is retained through *O*_Ali_.

The distance of maximum alignment *d*_Ali_ cancels in 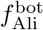 and therefore does not need to be specified for the robot. The kick length is *l* = v *τ* + *d*_*c*_, and the resulting target position is redrawn until it falls inside the disc of radius *R*.

### Why the interaction strengths differ from the simulation values

Since the robot leaves the angular functions un-normalized, a given *γ*_Att_ or *γ*_Ali_ produces a different turning amplitude in the two cases, and the values of Tables S2 and S3 should not be compared directly with those of Table S1. The peak values of the un-normalized functions used by the robot are max{*O*_Att_} = 1.05, 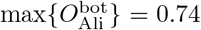 and 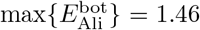, and *f*_Att_ reaches 2.87 *γ*_Att_ at *d* ≃ 0.23 m. The interaction ranges also differ, with *l*_Att_ = *l*_Ali_ = 0.20 m on the robot against 0.170 and 0.145 m in the simulations, as does the repulsion/attraction transition distance, 0.03 m against 0.015 m.

### Peak speed and kick duration

The kick duration is redrawn at every kick, with mean *τ*_avg_. Conversely, the peak speed used in the robot controller is not redrawn at every kick: with probability *p*_mem_ the robot reuses the speed measured by the tracking system, or the mean measured speed of its neighbors, and only with probability 1 − *p*_mem_ does it draw a new value. In Table S2, v_avg_ and v_min_ therefore parameterize that resampling law rather than the realized speed distribution, which for *p*_mem_ = 0.96 is dominated by the measured-speed feedback.

### Kick length and *τ*_0_

The robot advances the reference agent by *l* = v *τ* + *d*_*c*_ instead of integrating the exponential speed decay of Eq. (S3). The deceleration time *τ*_0_ therefore has no effect on the robot’s trajectory and is listed in Table S2 for completeness only. The additive term *d*_*c*_, between 7 and 10 mm, has no counterpart in the simulations: it guarantees a minimum displacement per kick, below which the position controller cannot resolve a new target.

### Effective radius

The rejection sampler confines the virtual reference agent to a disc of radius *R* between 0.19 and 0.225 m, smaller than the physical arena radius of 0.25 m. The margin absorbs the lag and overshoot of the tracking robot, which would otherwise contact the wall. When *n*_max_ = 50 consecutive proposals are rejected, the reference heading is reset to the local wall tangent plus an exponentially distributed offset, which unblocks agents trapped against the boundary.

### Lure-rescue routine

The lure is held to the robot magnetically and can occasionally detach, after which the robot would keep executing the model trajectory while the lure stays behind. The controller therefore monitors the distance between the tracked lure and the tracked robot, and when it exceeds *d*_rescue_ = 0.07 m it enters a recovery mode in which the model integration is suspended and the reference pose is instead set to the current position of the detached lure, so that the position controller drives the robot back to it. Normal model-driven motion resumes once the two are within 1 cm of each other. The routine allows the robot to re-attach the lure autonomously and avoids interrupting an experiment for manual intervention. It was enabled in the single-robot and pair conditions and disabled in the five-agent conditions.

### Condition-specific choices

Spontaneous noise was disabled (*γ*_R_ = 0) in both five-agent conditions, so that the robot’s heading was determined entirely by the wall and social terms. The *k* = 1 and *k* = 2 five-agent conditions used otherwise identical parameters: unlike in the simulations, *γ*_Att_ and *γ*_Ali_ were not re-tuned between them, so the comparison between *k* = 1 and *k* = 2 isolates the effect of the number of integrated neighbors at fixed interaction strengths. In the single-robot condition *k* = 0, which deactivates both social terms.

**Figure S1:**
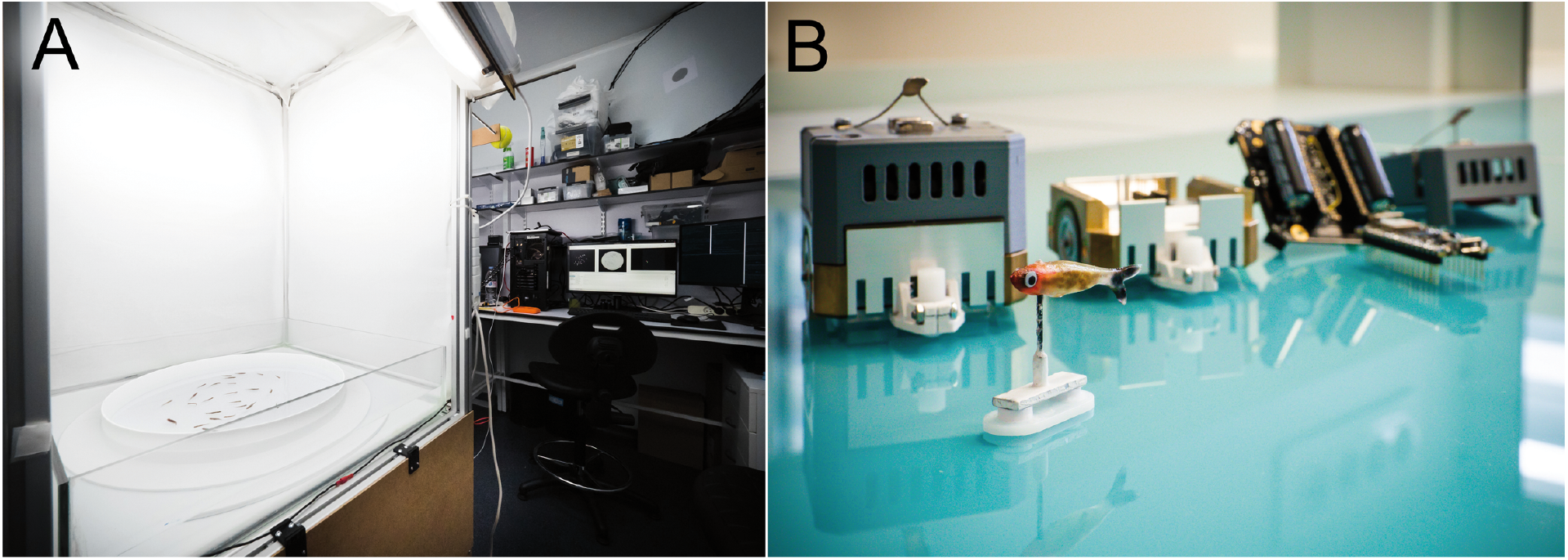
Experimental setup and biomimetic robotic lure. (A) Photograph of the experimental setup used for the closed-loop biohybrid experiments. (B) Photograph of the biomimetic fish lure and the Lurebot. Photographs by David Villa, ScienceImage, CBI, Toulouse.

**Figure S2:**
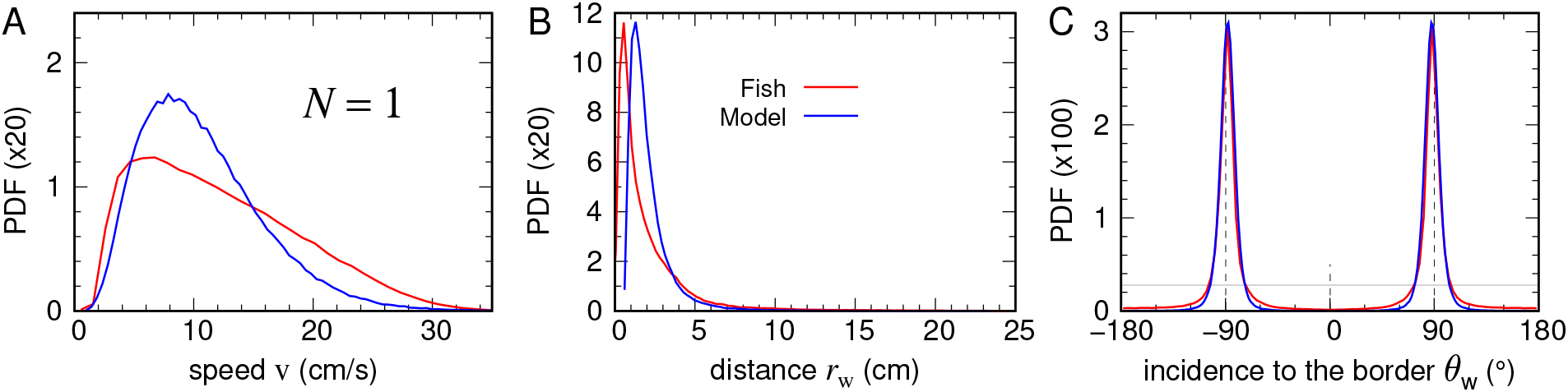
Individual behavioral descriptors for isolated fish. Probability density functions (PDFs) of (A) swimming speed v, (B) distance to the arena wall *r*w, (C) angle of incidence to the border *θ*w, measured in experiments with isolated fish (red) and corresponding numerical simulations (blue).

**Figure S3:**
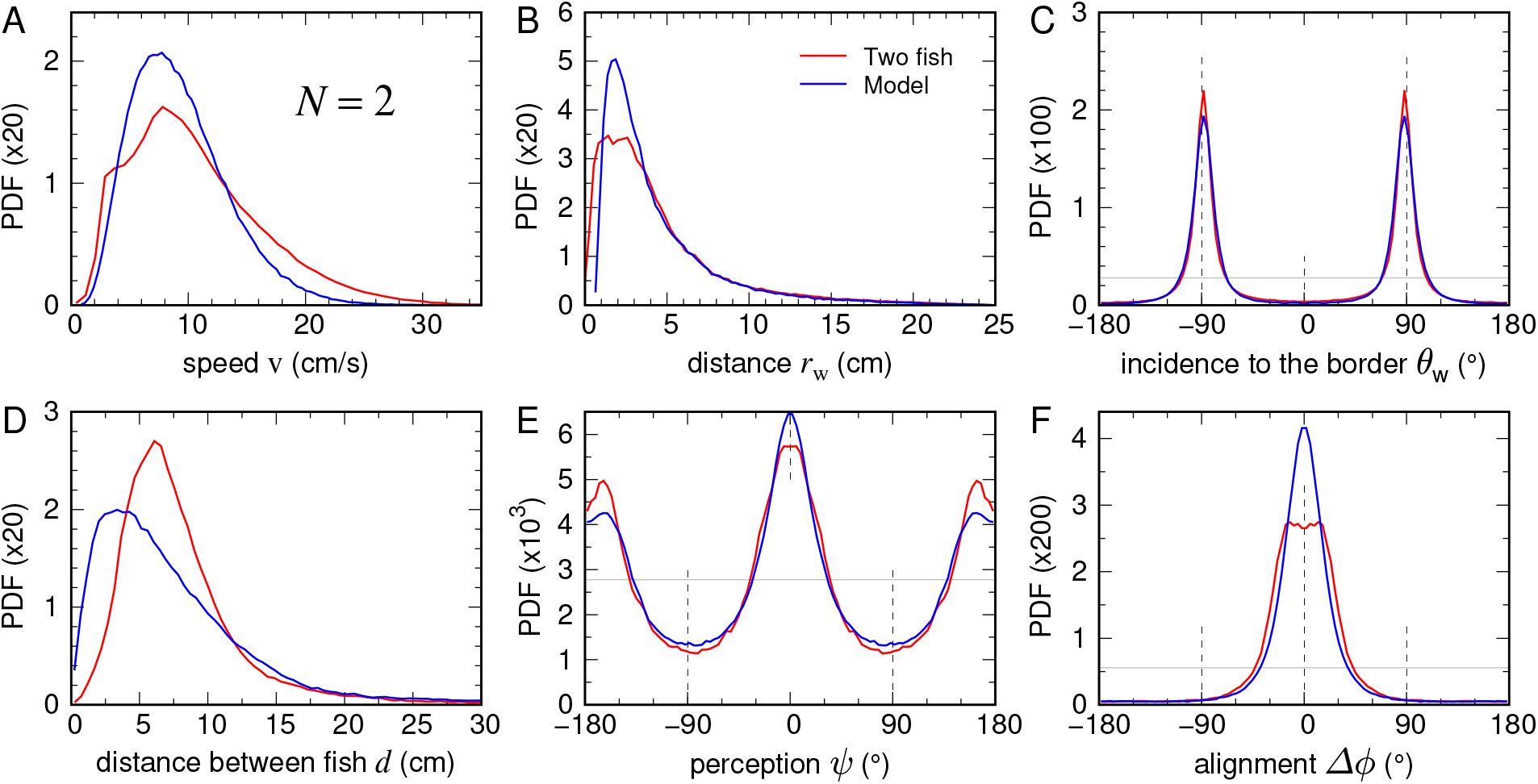
Pairwise behavioral descriptors in biological fish pairs. Probability density functions (PDFs) of (A) swimming speed v, (B) distance to the arena wall *r*_w_, (C) angle of incidence to the border *θ*_w_, (D) distance between fish *d*, (E) angle of perception *ψ*, and (F) relative alignment Δ*ϕ*, for biological fish pairs (red) and numerical simulations (blue).

**Figure S4:**
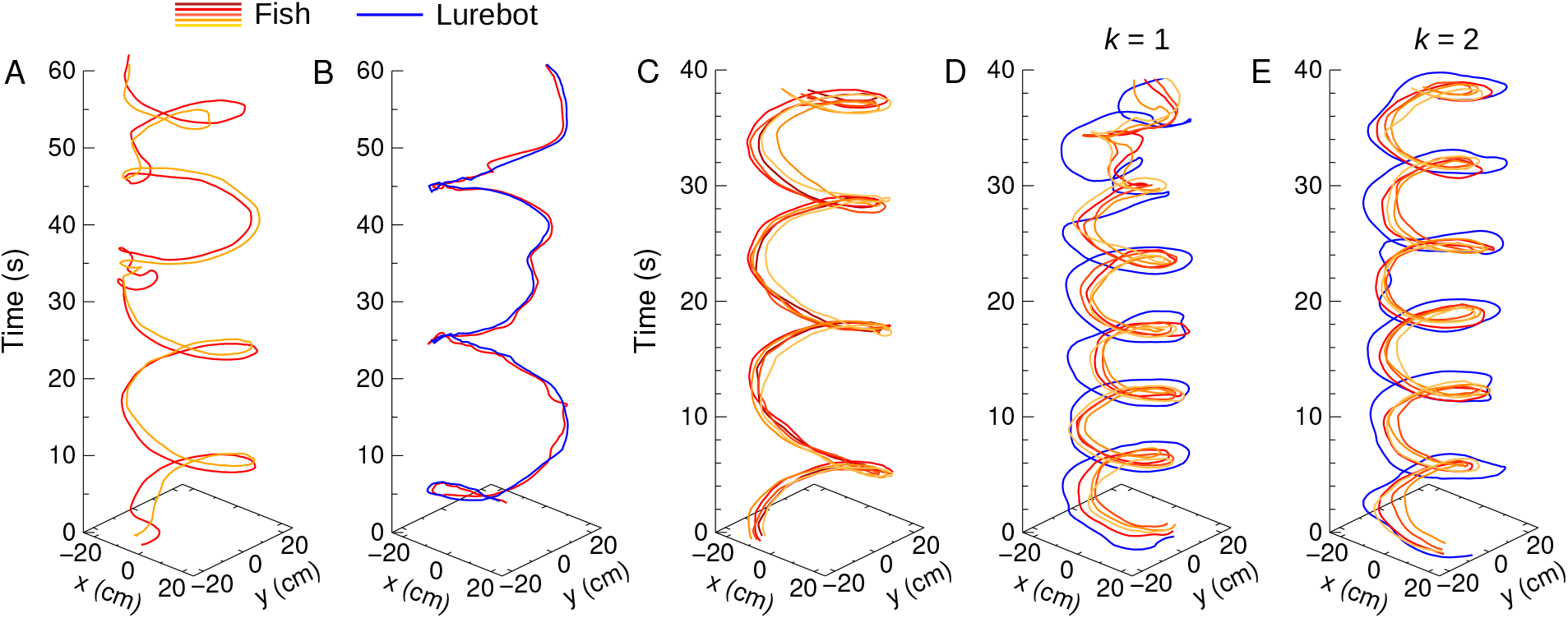
Representative trajectories in biohybrid groups of *N* = 2 and *N* = 5. Representative examples of individual trajectories of fish (orange-red shades) and lurebot (blue): (A) Two fish, (B) a fish and the lurebot, (C) five fish, (D) four fish and the lurebot interacting with its single most influential neighbor (*k* = 1), and (E) four fish and the lurebot interacting with its two most influential neighbors (*k* = 2).

**Figure S5:**
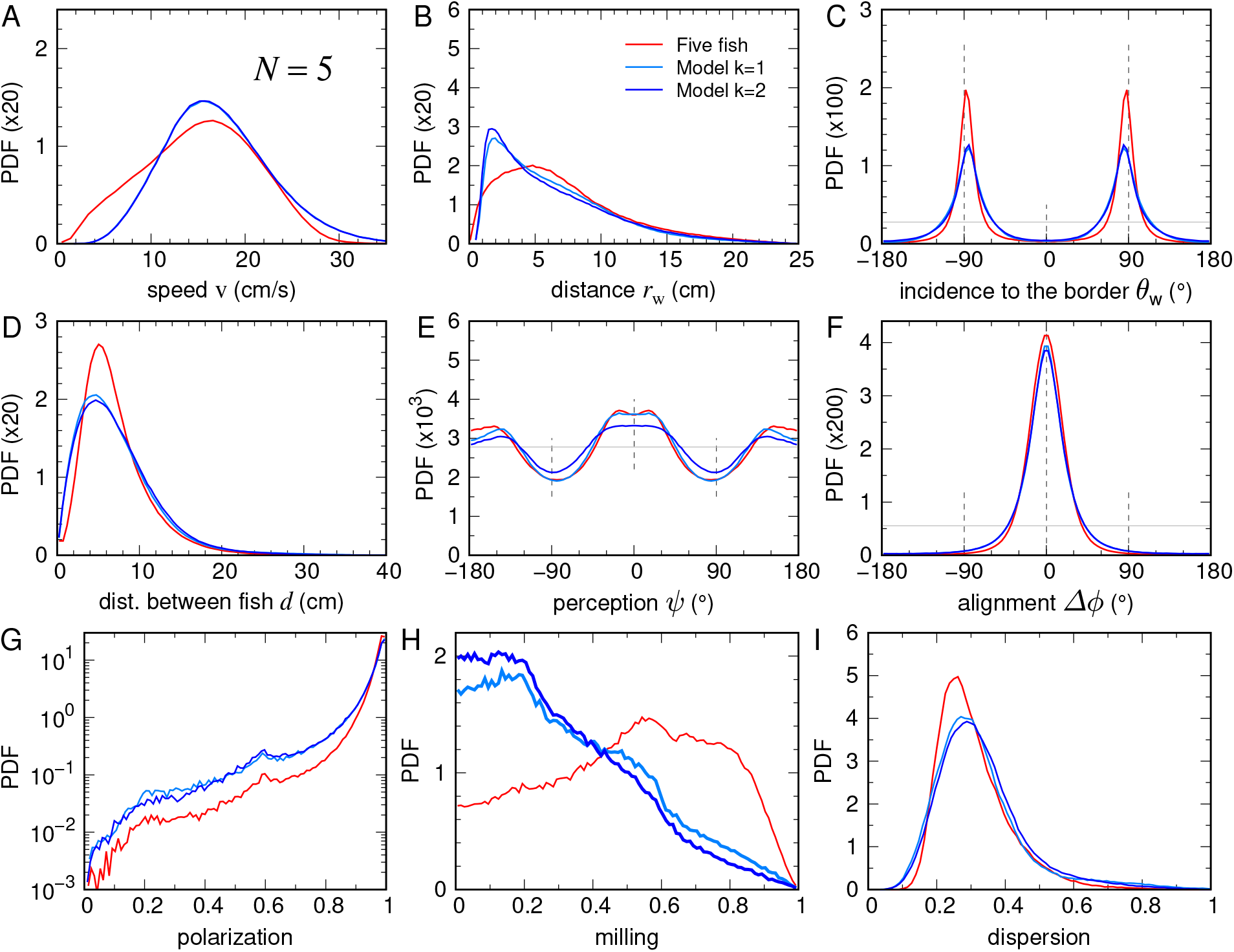
Behavioral descriptors in groups of five fish and numerical simulations. Probability density functions (PDFs) of (A) swimming speed v, (B) distance to the arena wall *r*_w_, (C) angle of incidence to the wall *θ*_w_, (D) distance between fish *d*, (E) viewing angle of perception *ψ*, (F) alignment Δ*ϕ*, (G) polarization, (H) milling, and (I) normalized spatial dispersion, for biological groups of five fish (red) and numerical simulations with *k* = 1 (light blue) and *k* = 2 (blue).

**Table S1:** Parameters of the behavioral model used in numerical simulations. Only the spontaneous noise amplitude (*γ*_R_), attraction strength (*γ*_Att_), and alignment strength (*γ*_Ali_) were adjusted independently for each experimental condition. All remaining parameters were kept fixed throughout the study.

| Parameter | Physical meaning | $N = 1$ | $N = 2$ | $N = 5, k = 1$ | $N = 5, k = 2$ |
| --- | --- | --- | --- | --- | --- |
| $\gamma_R$ | Spontaneous noise amplitude | 0.10 | 0.08 | 0.10 | 0.10 |
| $\gamma_{Att}$ | Pairwise attraction strength | – | 0.06 | 0.10 | 0.08 |
| $\gamma_{Ali}$ | Pairwise alignment strength | – | 0.025 | 0.070 | 0.045 |
| $v_{avg}$ (m/s) | Mean peak kick speed | 0.14 | 0.10 | 0.12 | 0.12 |
| $v_{min}$ (m/s) | Minimum peak kick speed | 0.04 | 0.04 | 0.13 | 0.13 |

| Parameter | Physical meaning | Value |
| --- | --- | --- |
| *Wall interaction* |  |  |
| $\gamma_w$ | Wall-repulsion strength | 0.50 |
| $l_w$ (m) | Wall interaction range | 0.06 |
| $\alpha$ | Noise attenuation near the wall | 0.67 |
| $\beta_w$ | Normalization constant of $O_w$ | 1.9612 |
| *Attraction–repulsion* |  |  |
| $d_{Att}$ (m) | Repulsion–attraction transition distance | 0.015 |
| $l_{Att}$ (m) | Attraction interaction range | 0.170 |
| $\beta_{Att}$ | Normalization constant of $O_{Att}$ | 1.43 |
| $\lambda_{Att}$ | Normalization constant of $E_{Att}$ | 0.90 |
| *Alignment* |  |  |
| $d_{Ali}$ (m) | Distance of maximum alignment | 0.015 |
| $l_{Ali}$ (m) | Alignment interaction range | 0.145 |
| $\beta_{Ali}$ | Normalization constant of $E_{Ali}$ | 0.90 |
| $\lambda_{Ali}$ | Normalization constant of $O_{Ali}$ | 1.43 |
| *Kick statistics* |  |  |
| $\tau_{avg}$ (s) | Mean kick duration | 0.55 |
| $\tau_0$ (s) | Deceleration time | 0.87 |
| *Geometry* |  |  |
| $R$ (m) | Arena radius | 0.25 |

**Table S2:**
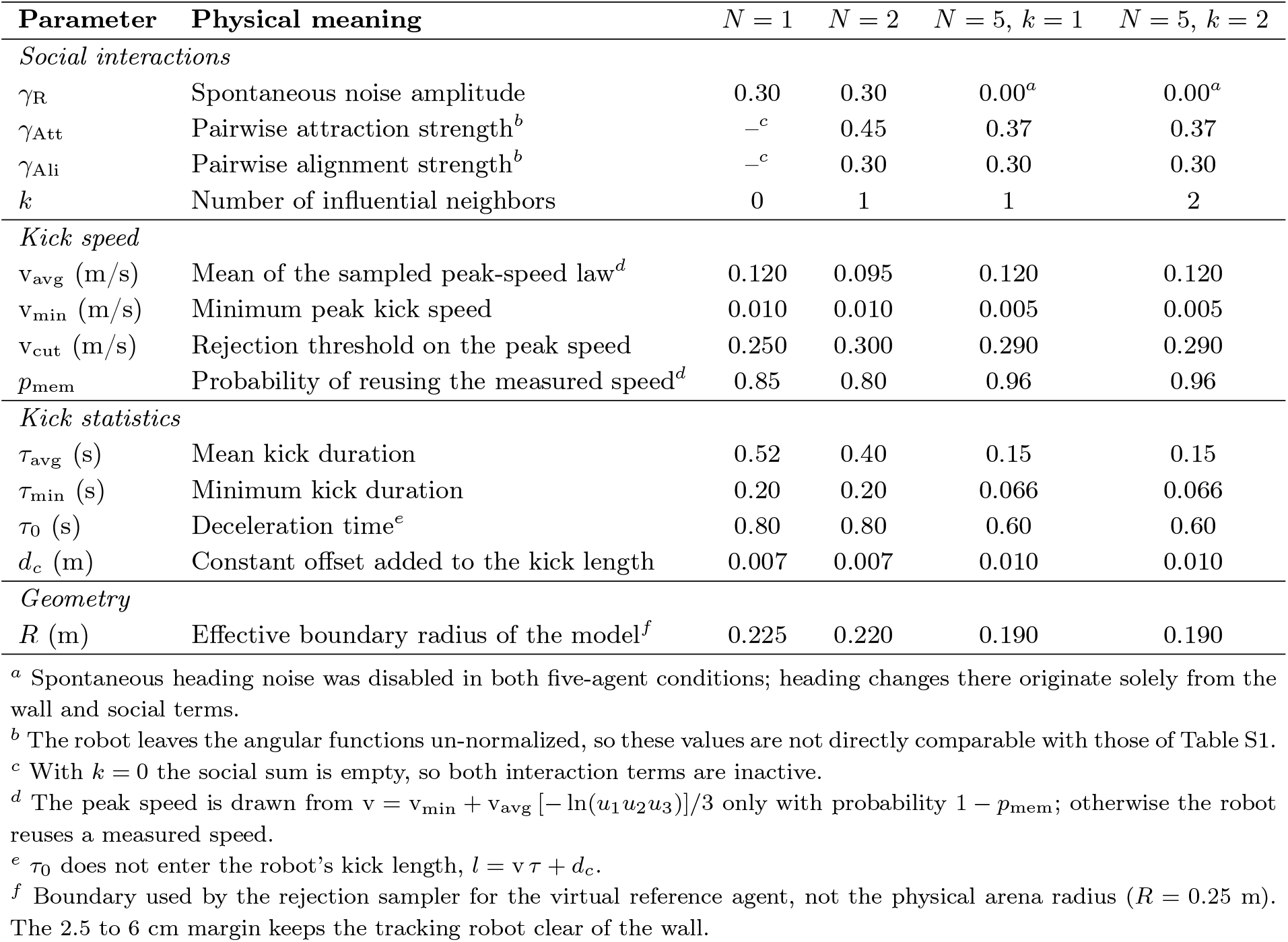
Condition-dependent parameters of the behavioral model running in the robot’s control loop. Unlike the numerical simulations, where only *γ*_R_, *γ*_Att_ and *γ*_Ali_ were adjusted per condition, the closed-loop experiments also required condition-specific kick-speed and kick-duration statistics and a condition-specific effective boundary radius, in order to accommodate the finite acceleration, tracking latency and physical extent of the robot. *k* denotes the number of most-influential neighbors actually integrated by the controller; it is a parameter of the robot model, not only a label of the experimental condition.

| Parameter | Physical meaning | $N = 1$ | $N = 2$ | $N = 5, k = 1$ | $N = 5, k = 2$ |
| --- | --- | --- | --- | --- | --- |
| *Social interactions* |  |  |  |  |  |
| $\gamma_R$ | Spontaneous noise amplitude | 0.30 | 0.30 | 0.00<sup>a</sup> | 0.00<sup>a</sup> |
| $\gamma_{Att}$ | Pairwise attraction strength<sup>b</sup> | —<sup>c</sup> | 0.45 | 0.37 | 0.37 |
| $\gamma_{Ali}$ | Pairwise alignment strength<sup>b</sup> | —<sup>c</sup> | 0.30 | 0.30 | 0.30 |
| $k$ | Number of influential neighbors | 0 | 1 | 1 | 2 |
| *Kick speed* |  |  |  |  |  |
| $v_{avg}$ (m/s) | Mean of the sampled peak-speed law<sup>d</sup> | 0.120 | 0.095 | 0.120 | 0.120 |
| $v_{min}$ (m/s) | Minimum peak kick speed | 0.010 | 0.010 | 0.005 | 0.005 |
| $v_{cut}$ (m/s) | Rejection threshold on the peak speed | 0.250 | 0.300 | 0.290 | 0.290 |
| $p_{mem}$ | Probability of reusing the measured speed<sup>d</sup> | 0.85 | 0.80 | 0.96 | 0.96 |
| *Kick statistics* |  |  |  |  |  |
| $\tau_{avg}$ (s) | Mean kick duration | 0.52 | 0.40 | 0.15 | 0.15 |
| $\tau_{min}$ (s) | Minimum kick duration | 0.20 | 0.20 | 0.066 | 0.066 |
| $\tau_0$ (s) | Deceleration time<sup>e</sup> | 0.80 | 0.80 | 0.60 | 0.60 |
| $d_c$ (m) | Constant offset added to the kick length | 0.007 | 0.007 | 0.010 | 0.010 |
| *Geometry* |  |  |  |  |  |
| $R$ (m) | Effective boundary radius of the model<sup>f</sup> | 0.225 | 0.220 | 0.190 | 0.190 |

<sup>a</sup> Spontaneous heading noise was disabled in both five-agent conditions; heading changes there originate solely from the wall and social terms.
<sup>b</sup> The robot leaves the angular functions un-normalized, so these values are not directly comparable with those of Table S1.
<sup>c</sup> With $k = 0$ the social sum is empty, so both interaction terms are inactive.
<sup>d</sup> The peak speed is drawn from $v = v_{min} + v_{avg} [-\ln(u_1 u_2 u_3)]/3$ only with probability $1 - p_{mem}$ ; otherwise the robot reuses a measured speed.
<sup>e</sup> $\tau_0$ does not enter the robot’s kick length, $l = v \tau + d_c$ .
<sup>f</sup> Boundary used by the rejection sampler for the virtual reference agent, not the physical arena radius ( $R = 0.25$ m). The 2.5 to 6 cm margin keeps the tracking robot clear of the wall.

**Table S3:** Parameters of the behavioral model held fixed across all robot experiments. None of these parameters was varied across conditions. The robot leaves the angular functions un-normalized, so all normalization constants are equal to 1.

| Parameter | Physical meaning | Value |
| --- | --- | --- |
| *Wall interaction* |  |  |
| $\gamma_w$ | Wall-repulsion strength | 0.23 |
| $l_w$ (m) | Wall interaction range | 0.06 |
| $\alpha$ | Noise attenuation near the wall | 0.60 |
| $\beta_w$ | Normalization constant of $O_w$ | 1 |
| *Attraction-repulsion* |  |  |
| $d_{Att}$ (m) | Repulsion-attraction transition distance | 0.030 |
| $l_{Att}$ (m) | Attraction interaction range | 0.200 |
| $\beta_{Att}$ | Normalization constant of $O_{Att}$ | 1 |
| $\lambda_{Att}$ | Normalization constant of $E_{Att}$ | $-(E_{Att}^{bot} \equiv 1)$ |
| *Alignment* |  |  |
| $d_{Ali}$ (m) | Distance of maximum alignment | $-^a$ |
| $l_{Ali}$ (m) | Alignment interaction range | 0.200 |
| $\lambda_{Ali}$ | Normalization constant of $O_{Ali}$ | 1 |
| $\beta_{Ali}$ | Normalization constant of $E_{Ali}$ | 1 |
| *Kick sampling* |  |  |
| $p_{mem,12}$ | Probability of adopting the neighbors' mean speed | 0.60 |
| $n_{max}$ | Rejected kicks before the heading is unblocked | 50 |

<sup>a</sup> $d_{Ali}$ cancels in $f_{Ali}^{bot} = (d_{Ali}/d) f_{Ali}$ and therefore does not need to be specified for the robot.

**Table S4:**
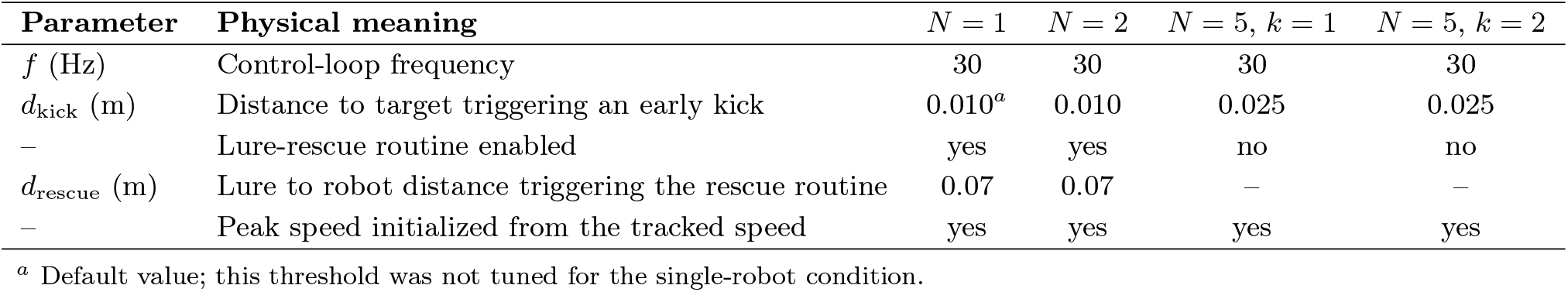
Control-loop and lure-tracking parameters. These parameters have no counterpart in the numerical simulations. They govern the closed-loop coupling between the model, which is integrated on a virtual reference agent, and the physical robot that tracks it.

| Parameter | Physical meaning | $N = 1$ | $N = 2$ | $N = 5, k = 1$ | $N = 5, k = 2$ |
| --- | --- | --- | --- | --- | --- |
| $f$ (Hz) | Control-loop frequency | 30 | 30 | 30 | 30 |
| $d_{kick}$ (m) | Distance to target triggering an early kick | $0.010^a$ | 0.010 | 0.025 | 0.025 |
| – | Lure-rescue routine enabled | yes | yes | no | no |
| $d_{rescue}$ (m) | Lure to robot distance triggering the rescue routine | 0.07 | 0.07 | – | – |
| – | Peak speed initialized from the tracked speed | yes | yes | yes | yes |

<sup>a</sup> Default value; this threshold was not tuned for the single-robot condition.

**Movie S1: Representative locomotor behavior of an isolated fish and the robotic fish**. The movie shows two representative 60-s recordings. The first sequence illustrates the spontaneous swimming behavior of an isolated *Hemigrammus rhodostomus* freely exploring the circular arena (radius = 25 cm). The second sequence shows the robotic fish (Lurebot) moving alone in the same arena under the control of the burst-and-coast behavioral model. The robotic fish reproduces the spontaneous locomotion, intermittent burst-and-coast swimming, and wall-following behavior characteristic of isolated fish in the absence of social interactions.

**Movie S2: Representative control and biohybrid pair experiments**. The movie presents two consecutive representative 60-s recordings. The first sequence shows a representative control experiment with two freely swimming *Hemigrammus rhodostomus* interacting in a circular arena (radius = 25 cm). The second sequence shows a representative biohybrid experiment in which one biological fish interacts with the autonomous robotic fish (Lurebot). The robotic fish continuously computes its burst-and-coast locomotion, wall avoidance, and social interactions from the real-time movements of the biological fish, establishing reciprocal closed-loop interactions throughout the experiment. The resulting coordinated swimming closely resembles that observed in control pairs of two biological fish.

**Movie S3: Representative control and biohybrid group experiments**. The movie consists of three consecutive 60-s sequences. The first sequence shows a representative control experiment with five freely swimming *Hemigrammus rhodostomus* in a circular arena (radius = 25 cm). The second sequence shows a representative biohybrid group composed of four biological fish and the autonomous robotic fish (Lurebot), in which the robot interacts exclusively with its single most influential neighbor (*k* = 1). The third sequence shows a representative biohybrid group under the same experimental conditions, except that the robotic fish simultaneously interacts with its two most influential neighbors (*k* = 2). In both biohybrid experiments, the robotic fish continuously updates its burst-and-coast locomotion, wall avoidance, and social interactions from the real-time movements of the biological fish, thereby establishing reciprocal closed-loop interactions throughout the experiment. Despite the different interaction hypotheses, both biohybrid groups exhibit cohesive and coordinated collective motion closely resembling that observed in the biological control group.

## References

Tamás Vicsek and Anna Zafeiris. Collective motion. Physics Reports, 517(3–4):71–140, 2012. doi: 10.1016/j.physrep.2012.03.004.

John E. Treherne and William A. Foster. Group transmission of predator avoidance behaviour in a marine insect: The trafalgar effect. Animal Behaviour, 29:911–917, 1981. doi: 10.1016/S0003-3472(81)80028-0.

Ashley J. W. Ward, James E. Herbert-Read, David J. T. Sumpter, and Jens Krause. Fast and accurate decisions through collective vigilance in fish shoals. Proceedings of the National Academy of Sciences, 108:2312–2315, 2011. doi: 10.1073/pnas.1007102108.

Daniel Grünbaum. Schooling as a strategy for taxis in a noisy environment. Evolutionary Ecology, 12: 503–522, 1998. doi: 10.1023/A:1006574607845.

David J. T. Sumpter. Collective Animal Behavior. Princeton University Press, Princeton, NJ, 2010.

Tao Xue, Xin Li, Guozheng Lin, Rodrigo Escobedo, Zhen Han, Xiaofei Chen, Clément Sire, and Guy Theraulaz. Tuning social interactions’ strength drives collective response to light intensity in schooling fish. PLoS Computational Biology, 19(11):e1011636, 2023. doi: 10.1371/journal.pcbi.1011636.

Guozheng Lin, Xin Li, Tao Xue, Rodrigo Escobedo, Zhen Han, Clément Sire, Vishwesha Guttal, and Guy Theraulaz. Experimental evidence of stress-induced critical state in schooling fish. PRX Life, 3: 033018, 2025. doi: 10.1103/PRXLife.3.033018.

Irene Giardina. Collective behavior in animal groups: theoretical models and empirical studies. HFSP Journal, 2(4):205–219, 2008. doi: 10.2976/1.2961038.

Ugo Lopez, Jacques Gautrais, Iain D. Couzin, and Guy Theraulaz. From behavioural analyses to models of collective motion in fish schools. Interface Focus, 2(6):693–707, 2012. doi: 10.1098/rsfs.2012.0033.

James E. Herbert-Read. Understanding how animal groups achieve coordinated movement. Journal of Experimental Biology, 219:2971–2983, 2016. doi: 10.1242/jeb.129411.

Nicholas T. Ouellette. A physics perspective on collective animal behavior. Physical Biology, 19:021004, 2022. doi: 10.1088/1478-3975/ac4805.

Ichiro Aoki. A simulation study on the schooling mechanism in fish. Bulletin of the Japanese Society of Scientific Fisheries, 48:1081–1088, 1982. doi: 10.2331/suisan.48.1081.

Akira Okubo. Dynamical aspects of animal grouping: swarms, schools, flocks, and herds. Advances in Biophysics, 22:1–94, 1986. doi: 10.1016/0065-227X(86)90003-1.

Andreas Huth and Christian Wissel. The simulation of the movement of fish schools. Journal of Theoretical Biology, 156:365–385, 1992. doi: 10.1016/S0022-5193(05)80681-2.

Andreas Huth and Christian Wissel. The simulation of fish schools in comparison with experimental data. Ecological Modelling, 75–76:135–146, 1994. doi: 10.1016/0304-3800(94)90013-2.

Hiro-Sato Niwa. Self-organizing dynamic model of fish schooling. Journal of Theoretical Biology, 171: 123–136, 1994. doi: 10.1006/jtbi.1994.1218.

Iain D. Couzin, Jens Krause, Richard James, Graeme D. Ruxton, and Nigel R. Franks. Collective memory and spatial sorting in animal groups. Journal of Theoretical Biology, 218:1–11, 2002. doi: 10.1006/jtbi.2002.3065.

Yael Katz, Kristoffer Tunström, Christos C. Ioannou, Cristián Huepe, and Iain D. Couzin. Inferring the structure and dynamics of interactions in schooling fish. Proceedings of the National Academy of Sciences, 108:18720–18725, 2011. doi: 10.1073/pnas.1107583108.

James E. Herbert-Read, Andrea Perna, Richard P. Mann, Timothy M. Schaerf, David J. T. Sumpter, and Ashley J. W. Ward. Inferring the rules of interaction of shoaling fish. Proceedings of the National Academy of Sciences, 108:18726–18731, 2011. doi: 10.1073/pnas.1109355108.

Jacques Gautrais, Francesco Ginelli, Richard Fournier, Sergio Blanco, Miguel Soria, Hugues Chaté, and Guy Theraulaz. Deciphering interactions in moving animal groups. PLoS Computational Biology, 8: e1002678, 2012. doi: 10.1371/journal.pcbi.1002678.

Daniel S. Calovi, Alexander Litchinko, Valentin Lecheval, Ugo Lopez, Alfonso Pérez-Escudero, Hugues Chaté, Clément Sire, and Guy Theraulaz. Disentangling and modeling interactions in fish with burst- and-coast swimming reveal distinct alignment and attraction behaviors. PLoS Computational Biology, 14(4):e1005933, 2018. doi: 10.1371/journal.pcbi.1005933.

Sarah B. Rosenthal, Colin R. Twomey, Andrew T. Hartnett, Henry S. Wu, and Iain D. Couzin. Revealing the hidden networks of interaction in mobile animal groups allows prediction of complex behavioral contagion. Proceedings of the National Academy of Sciences, 112(15):4690–4695, 2015. doi: 10.1073/pnas.1420068112.

Wouter Poel, Christian Winklmayr, and Pawel Romanczuk. Spatial structure and information transfer in visual networks. Frontiers in Physics, 9:716576, 2021. doi: 10.3389/fphy.2021.716576.

Liang Jiang, Luca Giuggioli, Andrea Perna, Rodrigo Escobedo, Valentin Lecheval, Clément Sire, Zhen Han, and Guy Theraulaz. Identifying influential neighbors in animal flocking. PLOS Computational Biology, 13(11):e1005822, 2017. doi: 10.1371/journal.pcbi.1005822.

Liang Lei, Rodrigo Escobedo, Clément Sire, and Guy Theraulaz. Computational and robotic modeling reveal parsimonious combinations of interactions between individuals in schooling fish. PLoS Computational Biology, 16(3):e1007194, 2020. doi: 10.1371/journal.pcbi.1007194.

Arnau Puy, Eduard Gimeno, Jordi Torrents, Pavel Bartashevich, Miguel C. Miguel, Romualdo Pastor-Satorras, and Pawel Romanczuk. Selective social interactions and speed-induced leadership in schooling fish. Proceedings of the National Academy of Sciences, 121(21):e2309733121, 2024. doi: 10.1073/pnas.2309733121.

Rotem Harpaz, Minh N. Nguyen, Armin Bahl, and Florian Engert. Precise visuomotor transformations underlying collective behavior in larval zebrafish. Nature Communications, 12:6578, 2021. doi: 10.1038/s41467-021-26808-7.

Yu Xiao, Xiang Lei, Zeyang Zheng, Yu Xiang, Yang-Yu Liu, and Xing Peng. Perception of motion salience shapes the emergence of collective motions. Nature Communications, 15:4779, 2024. doi: 10.1038/s41467-024-49026-0.

José Halloy Guillaume Sempo, Gilles Caprari, Colette Rivault, Mohsen Asadpour, Frédéric Tâche, Inès Saïd, Virginie Durier, Stéphanie Canonge, Jean-Marc Amé, Claire Detrain, Nikolaus Correll, Alcherio Martinoli, Francesco Mondada, Roland Siegwart, and Jean-Louis Deneubourg. Social integration of robots into groups of cockroaches to control self-organized choices. Science, 318(5853):1155–1158, 2007. doi: 10.1126/science.1144259.

José J. Faria, John R. G. Dyer, Romain O. Clément, Iain D. Couzin, Nicholas Holt, Ashley J. W. Ward, David Waters, and Jens Krause. A novel method for investigating the collective behaviour of fish introducing robofish. Behavioral Ecology and Sociobiology, 64:1211–1218, 2010. doi: 10.1007/s00265-010-0945-9.

Pasquale De Lellis, Elena Cadolini, Alessio Croce, Yizhou Yang, Mario Di Bernardo, and Maurizio Porfiri. Model-based feedback control of live zebrafish behavior via interaction with a robotic replica. IEEE Transactions on Robotics, 36(1):28–41, 2020. doi: 10.1109/TRO.2019.2943066.

Tim Landgraf, Georg H. W. Gebhardt, David Bierbach, Pawel Romanczuk, Lukas Musiolek, Verena V. Hafner, and Jens Krause. Animal-in-the-loop: Using interactive robotic conspecifics to study social behavior in animal groups. Annual Review of Control, Robotics, and Autonomous Systems, 4:487–507, 2021. doi: 10.1146/annurev-control-071920-095021.

Jens Krause, Alan F. T. Winfield, and Jean-Louis Deneubourg. Interactive robots in experimental biology. Trends in Ecology & Evolution, 26(7):369–375, 2011. doi: 10.1016/j.tree.2011.03.015.

Frank Bonnet, Alexey Gribovskiy, José Halloy and Francesco Mondada. Closed-loop interactions between a shoal of zebrafish and a group of robotic fish in a circular corridor. Swarm Intelligence, 12(3):227–244, 2018. doi: 10.1007/s11721-018-0153-6.

Vasileios Papaspyros, David Burnier, Rami Cherfan, Guy Theraulaz, Clément Sire, and Francesco Mon-dada. A biohybrid interaction framework for the integration of robots in animal societies. IEEE Access, 11:67640–67658, 2023. doi: 10.1109/ACCESS.2023.3291083.

Auke J. Ijspeert, Francesco Mondada, Emily Standen, and Guy Theraulaz. Swimming with robots: Investigating fish locomotion, sensing, and schooling behavior with robotic swimmers. Nature Communications, 17:5012, 2026. doi: 10.1038/s41467-026-72478-6.

Sachit Butail, Giacomo Polverino, Phong Phamduy, Francesco Del Sette, and Maurizio Porfiri. Fish-robot interactions in a free-swimming environment: Effects of speed and configuration of robots on live fish. In Proceedings of SPIE, volume 9083, page 90830R, 2014. doi: 10.1117/12.2044832.

Tim Landgraf, David Bierbach, Huy Nguyen, Niklas Muggelberg, Pawel Romanczuk, and Jens Krause. Robofish: Increased acceptance of interactive robotic fish with realistic eyes and natural motion patterns by live trinidadian guppies. Bioinspiration & Biomimetics, 11(1):015001, 2016. doi: 10.1088/1748-3190/11/1/015001.

Léo Cazenille, Bertrand Collignon, Yoann Chemtob, Frank Bonnet, Alexey Gribovskiy, Francesco Mon-dada, Nicolas Brédeche, and José Halloy. How mimetic should a robotic fish be to socially integrate into zebrafish groups? Bioinspiration & Biomimetics, 13(2):025001, 2018. doi: 10.1088/1748-3190/aaa8fd.

Vasileios Papaspyros, Frank Bonnet, Bertrand Collignon, and Francesco Mondada. Bidirectional inter-actions facilitate the integration of a robot into a shoal of zebrafish (Danio rerio). PLOS ONE, 14(8): e0220559, 2019. doi: 10.1371/journal.pone.0220559.

Vasileios Papaspyros, Guy Theraulaz, Clément Sire, and Francesco Mondada. Quantifying the biomimicry gap in biohybrid robot-fish pairs. Bioinspiration & Biomimetics, 19:046020, 2024. doi: 10.1088/1748-3190/ad4d4e.

Valentin Lecheval, Liang Jiang, Pierre Tichit, Clément Sire, Charlotte K. Hemelrijk, and Guy Theraulaz. Social conformity and propagation of information in collective U-turns of fish schools. Proceedings of the Royal Society B: Biological Sciences, 285:20180251, 2018. doi: 10.1098/rspb.2018.0251.

Maurizio Porfiri, Mert Karakaya, Raghu Ram Sattanapalle, and Sean D. Peterson. Emergence of in-line swimming patterns in zebrafish pairs. Flow, 1:E7, 2021. doi: 10.1017/flo.2021.5.

Diego A. Burbano-L. and Maurizio Porfiri. Modeling multi-sensory feedback control of zebrafish in a flow. PLOS Computational Biology, 17(1):e1008644, 2021. doi: 10.1371/journal.pcbi.1008644.

Scott Camazine, Jean-Louis Deneubourg, Nigel R. Franks, James Sneyd, Guy Theraulaz, and Eric Bonabeau. Self-Organization in Biological Systems. Princeton University Press, Princeton, NJ, 2001.

Iain D. Couzin. Collective cognition in animal groups. Trends in Cognitive Sciences, 13(1):36–43, 2009. doi: 10.1016/j.tics.2008.10.002.

Ramón Escobedo, Justine Reynaud, Stéphane Sanchez, Clément Sire, and Guy Theraulaz. Swimming speed of schooling fish controls social interaction strength in open-loop immersive virtual reality. Proceedings of the Royal Society B: Biological Sciences, 293(2072):20260146, 2026. doi: 10.1098/rspb.2026.0146. URL https://doi.org/10.1098/rspb.2026.0146.

L. Li, L. M. Chao, S. Wang, O. Deussen, and I. D. Couzin. Robotwin: A platform to study hydrodynamic interactions in schooling fish. IEEE Robotics & Automation Magazine, 31(1):10–17, 2024. doi: 10.1109/MRA.2023.3340776.

Patrick McMillen and Michael Levin. Collective intelligence: A unifying concept for integrating biology across scales and substrates. Communications Biology, 7:378, 2024. doi: 10.1038/s42003-024-06037-4.

Mehdi Moussaïd, Simon Garnier, Guy Theraulaz, and Dirk Helbing. Collective information processing and pattern formation in swarms, flocks, and crowds. Topics in Cognitive Science, 1(3):469–497, 2009. doi: 10.1111/j.1756-8765.2009.01028.x.

Francisco Romero-Ferrero, Matteo G. Bergomi, Robert C. Hinz, Francisco J. H. Heras, and Gonzalo G. de Polavieja. idtracker.ai: Tracking all individuals in small or large collectives of unmarked animals. Nature Methods, 16(2):179–182, 2019. doi: 10.1038/s41592-018-0295-5.

